# LRRK2 Phosphorylates Neuronal Elav RNA-Binding Proteins to Regulate Phenotypes Relevant to Parkinson’s Disease

**DOI:** 10.1101/2022.04.24.489327

**Authors:** Alyssa Pastic, Olanta Negeri, Aymeric Ravel-Chapuis, Alexandre Savard, My Tran Trung, Gareth Palidwor, Huishan Guo, Paul Marcogliese, James A. Taylor, Hideyuki Okano, Laura Trinkle-Mulcahy, Bernard J. Jasmin, David Park, Derrick Gibbings

## Abstract

Parkinson’s disease (PD) is characterized by accumulation of *α*-synuclein and the loss of dopaminergic neurons. Mutations which cause an increase in the kinase activity of Leucine-Rich-Repeat Kinase-2 (LRRK2) are a major inherited cause of PD. Research continues to determine which targets LRRK2 phosphorylates to cause disease. Polymorphisms in the locus of *ELAVL4,* an RNA-binding protein are a risk-factor for Parkinson’s disease and an *ELAV* family member was identified in *Drosophila* as required for pathology instigated by human mutant LRRK2. We discovered that three neuronal ELAVs including ELAVL4 (also known as HuD) are phosphorylated by LRRK2. This controls binding of neuronal ELAVs to mRNA and their post-transcriptional regulation of mRNA abundance and splicing in neuronal cell lines and the mouse midbrain. LRRK2 G2019S functionally inhibits neuronal ELAVs effects on mRNA abundance, while enhancing their effects on mRNA splicing. The combination of *LRRK2 G2019S* and *ELAVL4^-/-^* causes accumulation of LRRK2 and *α*-synuclein, loss of dopaminergic neurons and motor deficits. Targets of neuronal ELAVs are also selectively misregulated in cells and tissues of PD patients. Together, this suggests that misregulation of neuronal ELAVs, triggered by LRRK2 mutations may contribute to the characteristic pathology of Parkinson’s disease.

**Brief Summary:** LRRK2, a kinase linked to Parkinson’s disease, phosphorylates the neuronal ELAV RNA-binding proteins to aggravate key hallmarks of Parkinson’s disease including accumulation of *α*-synuclein and motor deficits in mice.

## Introduction

Parkinson’s disease is a progressive neurodegenerative disease, with both idiopathic and genetic causes. Pathology in patients with PD is primarily characterized by accumulation of *α*-synuclein and the progressive loss of dopaminergic neurons in the substantia nigra of the midbrain which leads to characteristic tremors and bradykinesia^1^.

Mutations in several genes can cause forms of PD^2, 3^. Duplication and triplication of the *α-synuclein* gene, *SNCA*^4, 5^, or variants which affect its expression^6^, can cause PD. Over-expression of *α*-synuclein elicits similar Parkinson’s-like symptoms in mice^7^, emphasizing the critical role of *α*-synuclein levels in PD. The most common cause of familial PD are mutations in *Leucine-Rich-Repeat Kinase 2* (*LRRK2*)^8, 9^. Several mutations in *LRRK2* that cause PD result in over-active kinase activity, including the most common mutation G2019S^10^. This suggests that target proteins phosphorylated by LRRK2 are the direct causes of patient pathology and symptoms. LRRK2 can phosphorylate proteins regulating translation including the small ribosomal subunit 15^11^, Argonautes^12^ and eIF4EBP^13^ but there is debate about whether these are physiological substrates of LRRK2. Recent unbiased phosphoproteomic studies have highlighted that LRRK2 phosphorylates RAB proteins including RAB10, RAB8a/b, RAB35, Rab12 and RAB29 to govern organelle dynamics and ciliogenesis^14–16^ by locking Rabs in a GTP-bound form. Other targets of LRRK2 kinase include WAVE2^17^ and endophilin A which also regulate membrane dynamics^18^. Phosphorylation of targets including Rab10 and other Rabs^19, 20^ and WAVE2^17^ have been validated in patient cells. Phosphorylation of RABs by LRRK2 may mediate effects ascribed to LRRK2 at the cellular level including effects on autophagy^21, 22^, lysosomes, vesicular trafficking^23^ and ciliogenesis^24^. While phosphorylated Rabs are readily detected in cell lines like HEK293 they are at low or undetectable levels in the brain^19, 25^, potentially due to the abundance of the relevant phosphatase^26^. Indeed, phosphorylation of Rab10 pThr73 was not dependent on Lrrk2 in the brain, while it was in peripheral organs^27^. Even in cells and peripheral tissues the phosphorylated Rabs only account for about 1% of total cellular Rabs^14, 26^. The impact of Rab phosphorylation by Lrrk2 is suggested by studies in non-mammals including *Drosophila*^13, 17, 28^ but their impact on crucial phenotypes in Parkinson’s disease like loss of dopaminergic neurons, accumulation of alpha-synuclein and motor deficits is unresolved.

Large-scale efforts to discover Lrrk2 substrates have mostly used generic cell lines like mouse embryonic fibroblasts rather than neurons or brain tissues. This leaves many potential brain-specific targets of Lrrk2 left to be discovered. Interestingly, a previous genome-wide screen in *Drosophila* to identify proteins required to induce pathology induced by PD-linked mutations in LRRK2 identified an RNA-binding protein of the *ELAV* (*Embryonic Lethal Abnormal Vision)* family^28^. Notably, studies in multiple independent patient cohorts found that single-nucleotide polymorphisms in the region of an *ELAV* family member, *ELAVL4,* are linked with susceptibility to or age-of-onset of PD^29–31^. This suggests that *ELAVL4* may impact PD pathogenesis and may control LRRK2-dependent pathology. The mechanisms by which *ELAVL4* may be regulated in PD or impact processes or phenotypes associated with PD is not known.

ELAVL4 is more commonly known as HuD and is expressed nearly exclusively in neurons^32^. In mice and humans, HuD has two homologues which are also preferentially expressed in neurons called HuB and HuC (or ELAVL2 and ELAVL3). HuB, HuC and HuD bind U-rich motifs in mRNAs in the 3’UTR, 5’UTR and introns of mRNAs^33, 34^. HuD contains three RNA-Recognition-Motifs (RRM). RRM1 and -2 bind to U-rich RNA motifs, while RRM3 binds to the polyA tail of mRNA^32, 35, 36^. By binding in introns, HuC and HuD have important impacts on splicing of mRNAs to which they bind^33^. In addition, HuC and HuD can either stabilize^37–40^ or destabilize mRNAs or regulate their translation^41–44^. By binding to hundreds of mRNAs and controlling their splicing, stability and translation HuB, -C and -D broadly regulate cellular processes^33, 34, 37, 45, 46^. mRNAs bound to and regulated by HuD include several involved in neuronal survival and formation of synaptic connections like *Brain-Derived Neurotrophic Factor* (*BDNF*)^47^. Neuronal cells lacking HuD have impaired dendritic outgrowth^47–49^. Mice lacking HuD exhibit motor deficits such as susceptibility to epileptic attacks and hind-limb clasping phenotypes^33, 48^. The impact of loss of HuD on Parkinson’s-like pathology and symptoms has not been previously investigated in detail, so it remains unclear to what extent HuB, HuC and HuD could contribute to Parkinson’s disease.

Here, we demonstrate that LRRK2 phosphorylates HuB, C and D, regulating their binding to mRNA, and inhibiting their post-transcriptional control of expression and splicing of mRNAs in neuronal cells and the mouse brain. When mice are sensitized by loss of *HuD* the hyperactive *LRRK2 G2019S* phosphorylates HuB and HuC and exacerbates hallmarks of PD including accumulation of *α*-synuclein, loss of dopaminergic neurons and motor deficits.

## Results

### LRRK2 Controls Binding of Neuronal ELAVs to Target mRNAs

We selected the human neuroblastoma cell line SH-SY5Y to study whether LRRK2 and its kinase activity affect HuD function. Flag-HuD was immunoprecipitated (Fig.1a) and bound mRNAs were quantified by RT-qPCR (Fig.1b). mRNAs which do not bind HuD, like *GAPDH* mRNA, were not enriched in HuD immunoprecipitates (Fig.1b). In contrast, mRNAs known to bind HuD including *BDNF, HuD* and *p21* were enriched by 10- to 1000-fold in HuD immunoprecipitates compared to levels in immunoprecipitates of control antibody (Fig.1b)^37, 50^. This validates against previous literature that the immunoprecipitation strategy used to isolate HuD accurately detects HuD binding to mRNAs. *SNCA* mRNA has motifs similar to those required for binding to neuronal ELAVs^34^ and was enriched in HuD immunoprecipitates as was the mRNA encoding LRRK2 (Fig.1b). The 3’UTRs of *Scna and Lrrk2* mRNAs also contain several canonical binding sites for neuronal ELAVs (Supplementary Dataset 1). This suggests these mRNAs encoding proteins highly relevant to PD may also be regulated by HuD.

**Figure 1.**
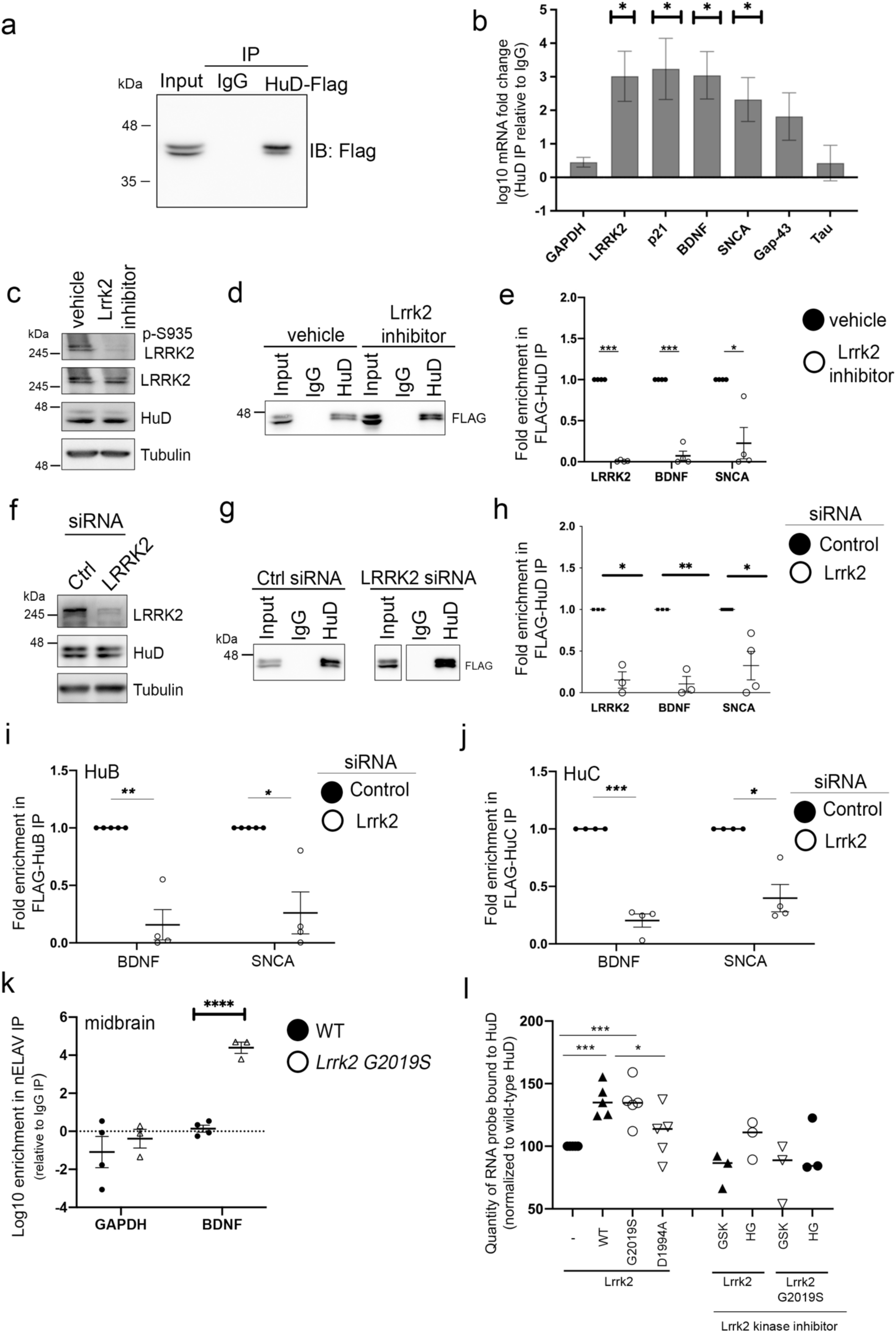
LRRK2 Kinase Activity Controls Binding of Neuronal ELAV Proteins to mRNA. **(a)** Western blot of immunoprecipitates of anti-Flag antibody or an isotype control antibody (IgG) from cells transfected with plasmid expressing Flag-HuD. **(b)** Log10 enrichment of the indicated mRNAs (x-axis) in immunoprecipitates of Flag-HuD from SH-SY5Y cells normalized to levels in IgG control immunoprecipitates measured by RT-qPCR in 4-6 independent experiments. * p<0.05, ** p<0.01, *** p<0.001 ANOVA with Tukey’s correction where multiple groups were compared to IgG control. **(c)** Western blot of LRRK2 phosphoserine-935 in cell lysates after treatment with control vehicle or GSK2578215A LRRK2 kinase inhibitor. **(d)** Western blot of Flag in immunoprecipitates of Flag-HuD from cells treated with control vehicle or GSK2578215A LRRK2 kinase inhibitor. **(e)** Quantification of RNAs by RT-qPCR in Flag-HuD immunoprecipitates from SH-SY5Y cells over four independent experiments from cells treated with control vehicle or GSK2578215A LRRK2 kinase inhibitor. **(f)** Western blot of LRRK2 in cell lysates after treatment with control siRNA or LRRK2 siRNA. **(g)** Western blot of Flag in immunoprecipitates of Flag-HuD from cells treated with control siRNA or LRRK2 siRNA. **(h)** Quantification of RNAs by RT-qPCR in Flag-HuD immunoprecipitates from SH-SY5Y cells treated with control siRNA or LRRK2 siRNA over 3-4 independent experiments. **(i,j)** Quantification of RNAs by RT-qPCR in immunoprecipitates of Flag-HuB **(i)** or Flag-HuC **(j)** from SH-SY5Y cells treated with control siRNA or LRRK2 siRNA over 2-4 independent experiments. **(k)** Quantification of RNAs by RT-qPCR in neuronal ELAV immunoprecipitates from midbrain of 3-4 wild-type and *LRRK2 G2019S* mice. **(l)** Quantification of gel-shift assay of HuD binding to *γ*-32P labeled U-rich RNA containing neuronal ELAV binding sites from *BDNF* 3’UTR or a control RNA from another segment of the *BDNF* 3’UTR. The effect of the presence of LRRK2 variants and LRRK2 kinase inhibitors (GSK2578215A [GSK], or HG-10-102-01 [HG]) was tested. * p<0.05, ** p<0.01, *** p<0.001 ANOVA with Tukey’s correction where multiple groups were compared, one-sample t-test (in e,h,I,j where one group was compared.

We tested whether LRRK2 affected HuD binding to mRNA. In cells treated with a LRRK2 kinase inhibitor, phosphorylation of LRRK2 at S935 was decreased as expected (Fig.1c). In Flag-HuD immunoprecipitates, mRNA encoding BDNF, SNCA, and LRRK2 detected by RT-qPCR was decreased when LRRK2 kinase was inhibited with GSK2578215A (Fig.1d,e, Supplementary Fig.1a). To ensure this was not due to an off-target effect of the kinase inhibitor, a similar experiment was performed after knocking down *LRRK2* with siRNA (Fig.1f). Silencing of *LRRK2* with siRNA (Fig.1f) decreased binding of HuD to *BDNF*, *SNCA*, and *LRRK2* mRNAs (Fig.1g,h). Binding of the two conserved HuD homologues, HuB and HuC, to *BDNF* and *SNCA* mRNAs was also reduced by *LRRK2* siRNA (Fig.1i,j, Supplementary Fig.1b) or LRRK2 kinase inhibitor (Supplementary Fig.1c-f). To evaluate the relevance of this to PD, similar experiments were performed in midbrain homogenate from wild-type and *LRRK2 G2019S* mice. Available antibodies cannot differentiate the highly homologous neuronal ELAVs (HuB, HuC, and HuD)^33^. Neuronal Elavs bound increased amounts of *Bdnf* in midbrain of mice expressing the hyperactive kinase *LRRK2 G2019S* compared to wild-type controls, while binding to non-specific *Gapdh* mRNA was not impacted (Fig.1k, Supplementary Fig.1g).

Results above demonstrate that depleting LRRK2 or inhibiting its kinase activity, reduced binding of HuD to several of its target mRNAs in cells, while increasing LRRK2 kinase activity in mouse brain increased neuronal ELAV binding to mRNA (Fig.1a-k). These effects on HuD could be due to indirect effects of LRRK2 on other cellular processes or kinases. To test whether LRRK2 directly affects binding of HuD to RNA, we utilized gel shift assays with purified HuD (Supplementary Fig.2a). HuD preferentially binds U-rich motifs ^33, 37^, including one in the *BDNF* 3’UTR, as previously shown by others in gel-shift assays^47^ (Supplementary Fig.2b). HuD, but not the control GST protein, bound to synthetic RNA containing this site, whereas no binding was detected to RNA derived from another site in the *BDNF* 3’UTR (Supplementary Fig.2c). This validates that in this assay HuD specifically binds to RNAs containing its defined motifs^33, 37, 47^. No binding of LRRK2 to these RNAs was detected (Supplementary Fig.2c). Equal amounts of wild-type LRRK2, kinase-dead LRRK2 D1994A mutant, or over-active kinase mutant LRRK2 G2019S were incubated with HuD and its target RNA. Interestingly, LRRK2 or LRRK2 G2019S increased HuD binding to RNA; an effect prevented by either of two independent LRRK2 kinase inhibitors (Figure 1l, Supplementary Fig.2c). Kinase-dead LRRK2 D1994A did not affect binding of HuD to RNA (Figure 1l). This demonstrates that LRRK2 directly increases binding of HuD to its U-rich target RNAs and this depends on the kinase activity of LRRK2. Collectively, this demonstrates that the kinase activity of LRRK2 promotes binding of HuD to target mRNAs in three contexts: *in vitro* with recombinant proteins, in cell cultures and in the mouse midbrain.

### LRRK2 Phosphorylates Neuronal ELAVs

Because LRRK2 effects on HuD binding to target mRNAs was controlled by its kinase activity we hypothesized that LRRK2 phosphorylates HuD. In intact cells, a Proximity Ligation Assay (PLA) detected LRRK2 in proximity to HuD (Fig.2a), excluding a post-lysis artefact. siRNA targeting *HuD* or *LRRK2* reduced these PLA signals, validating that HuD and LRRK2 were specifically detected in proximity to each other (Fig.2a,b). LRRK2 was also retrieved with HuD by immunoprecipitation (Fig.2c). This demonstrates that LRRK2 is positioned to phosphorylate HuD.

**Figure 2.**
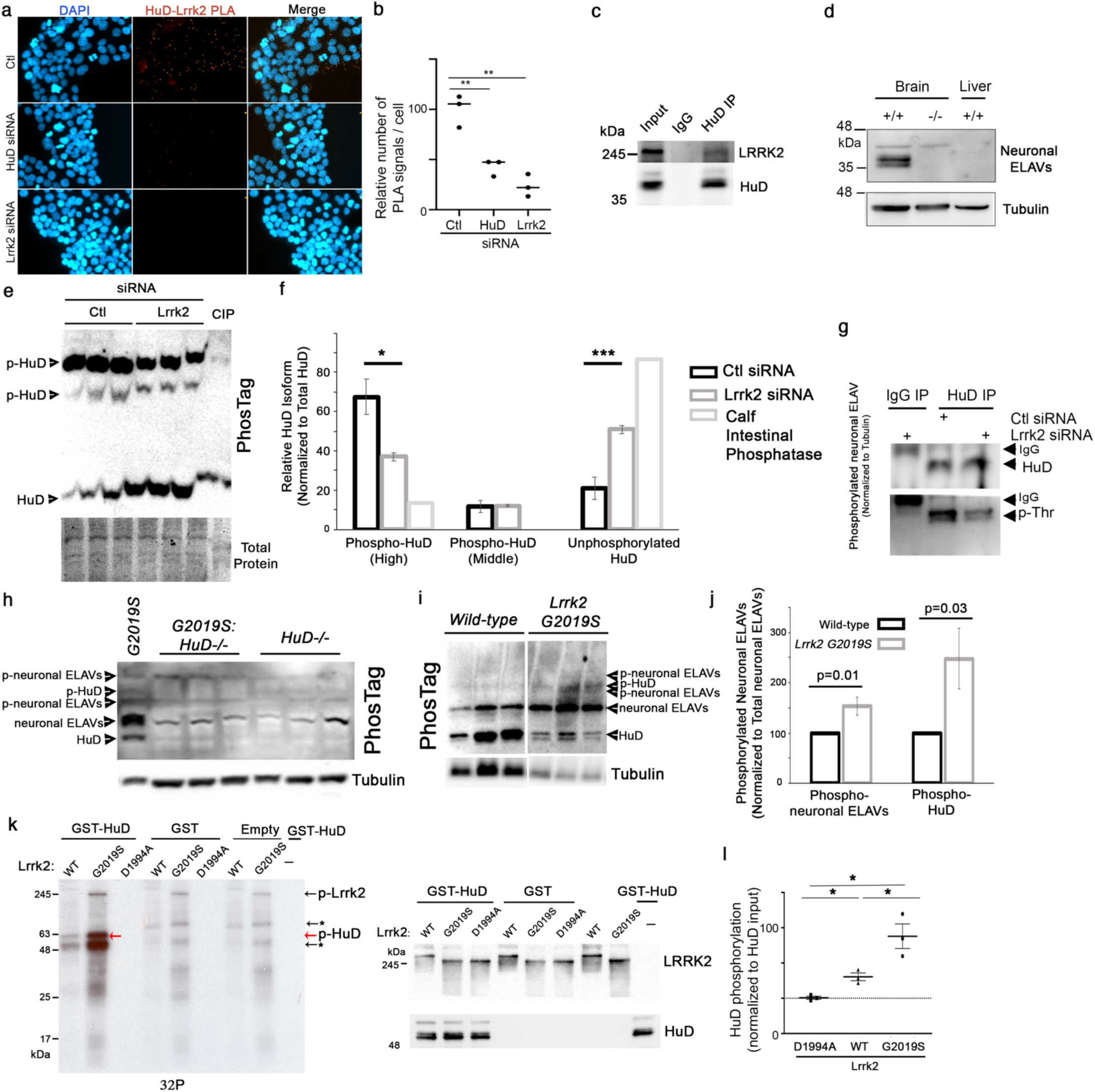
LRRK2 Phosphorylates Neuronal ELAV Proteins *in vitro*. **(a)** Representative images of proximity ligation assays between HuD and LRRK2 in SH-SY5Y cells treated with control siRNA, HuD siRNA or LRRK2 siRNA. **(b)** Quantification of proximity ligation assays as shown in (b). ** p<0.01 (t-test). **(c)** Western blot of LRRK2 and HuD in immunoprecipitates of endogenous HuD from Neuro2a cells. **(d)** Western blot of neuronal ELAVs in brain and liver of wild-type and *HuD^-/-^* mice. **(e)** Representative Western blot of PhosTag gels for Flag-HuD (anti-Flag) in SH-SY5Y cells treated with control or Lrrk2 siRNA, or where lysates were treated with CIP (Calf Intestinal Phosphatase) to dephosphorylate HuD. **(f)** Quantification of phosphorylated HuD in PhosTag blots of lysates of SH-SY5Y cells in (d). **(g)** Western blot of phospho-Threonine on Flag-HuD immunoprecipitates from SH-SY5Y cells treated with control siRNA or siRNA targeting LRRK2. **(h)** Representative Western blot of PhosTag gels for neuronal ELAVs in midbrain of *LRRK2 G2019S*, *HuD-^/-^* and *LRRK2 G2019S*: *HuD-^/-^* mice, showing specificity of bands for HuD. **(i)** Representative Western blot of PhosTag gels for neuronal ELAVs in midbrain of wild-type and *LRRK2 G2019S* mice, **(j)** Quantification of phosphorylated HuD and phosphorylated neuronal ELAVs in PhosTag blots of lysates of midbrain of wild-type (n=4) and *LRRK2 G2019S* (n=4) mice. **(k)** Autoradiographic exposure of SDS- PAGE gel after incubation of recombinant GST-HuD or controls with LRRK2 WT, LRRK2 G2019S and LRRK2 D1994A in the presence of *γ*-32P. Red arrow highlights phosphorylated HuD. Black arrows highlight autophosphorylated LRRK2, fragments of LRRK2, or contaminants are also observed by Coomassie total protein stain (Supplementary 2a). Below, Western blot of HuD and LRRK2 in samples from autoradiographic gels above. **(l)** Quantification of HuD phosphorylation on *γ*-32P gels as in (i) over n=3 independent experiments. *γ*-32P at the mass of GST-HuD was normalized to HuD detected by western blot in each experiment **(i)**. * p<0.05, ** p<0.01, *** p<0.001 ANOVA with Tukey’s correction where multiple groups were compared, t-test (two tailed assuming unequal variance) where one group was compared.

Available antibodies detect all of the highly homologous neuronal ELAVs (HuB, HuC and HuD) at slightly different masses. As expected, in mouse midbrain, an antibody recognizing neuronal ELAVs detected multiple bands for HuB, -C and -D some of which were absent in brain of *HuD-/-* mice and liver which does not express neuronal ELAVs, validating these as HuD and neuronal ELAVs respectively (Fig.2d). To assess the impact of LRRK2 on phosphorylation of HuD, we utilized PhosTag gels which cause a mass-shift in phosphorylated proteins (Fig.2e,f).Two higher mass bands of Flag-HuD were detected on PhosTag gels, and these were nearly eliminated by dephosphorylation with Calf Intestinal Phosphatase (CIP), confirming their phosphorylation (Fig.2e,f). Unphosphorylated HuD was detected at the bottom of the gel. Resembling the effect of CIP, treatment of cells with Lrrk2 siRNA reduced phosphorylation of the one of the phospho-HuD species and caused an accumulation of unphosphorylated HuD. To confirm with a second independent method that LRRK2 controls phosphorylation of HuD, an antibody recognizing phospho-Threonine was used. In the dopaminergic SH-SY5Y cell line HuD was detected anti- phospho-Threonine in immunoprecipitates, and this signal was reduced when LRRK2 was silenced with siRNA (Fig.2g). Some phosphorylation of HuD on Threonines was still detected after depletion of LRRK2 potentially due to incomplete depletion of LRRK2 by siRNA, or the phosphorylation of HuD by other kinases in addition to LRRK2.

To assess the impact of LRRK2 on phosphorylation of neuronal ELAVs in mice, we utilized PhosTag gels (Fig.2h). Several mass-shifted bands were detected with antibody recognizing neuronal ELAVs in knock-in *LRRK2 G2019S* mice. These mass-shifted bands were eliminated when lysates were dephosphorylated with CIP, validating these as *bona fide* phosphorylation events on neuronal ELAVs (Supplementary Fig.3a). In *HuD^-/-^* and *LRRK2 G2019S:HuD^-/-^* mice one unphosphorylated band and one mass-shifted band were absent, validating these as unphosphorylated and phosphorylated HuD respectively (Fig.2h). An additional two mass-shifted bands, presumably phosphorylated versions of HuB and/or HuC, were also detected at higher mass with anti-neuronal ELAV antibodies (Fig.2h). Having validated the specificity of the PhosTag analysis for HuD and *bona fide* phosphorylation, we quantified the effect of *LRRK2 G2019S* on phosphorylation of neuronal ELAVs. The phosphorylation of HuD was increased over 2-fold in midbrain of *LRRK2 G2019S* mice compared to wild-type littermates (Fig.2i,j) consistent with 2-3-fold increase in kinase activity of LRRK2 G2019S vs. wild- type LRRK2^10^. The phosphorylation of a band corresponding to other neuronal ELAVs was also significantly increased in *LRRK2 G2019S* compared to wild-type controls (Fig.2i,j).

The evidence above demonstrates that HuD and other neuronal ELAVs are phosphorylated in a LRRK2- dependent manner in relevant cell lines and the mouse midbrain. To test whether LRRK2 can directly phosphorylate neuronal ELAVs we utilized *in vitro* assays using recombinant wild-type LRRK2, the Parkinson’s-linked hyperactive variant LRRK2 G2019S and the kinase-dead LRRK2 variant D1994A (Supplementary Fig.2a). LRRK2 autophosphorylation was observed, as expected, with equal amounts of wild-type LRRK2 and LRRK2 G2019S, but not the D1994A kinase-dead mutant (black arrows, Fig.2k). LRRK2 G2019S exhibited increased autophosphorylation compared to wild-type LRRK2 consistent with its hyperactive kinase activity (Fig.2k,l). Phosphorylation of some bands at sizes lower than full-length Lrrk2 in preparations of purified Lrrk2 were observed. While these could be contaminating proteins, they are likely autophosphorylated fragments of Lrrk2 (asterisks, see “Empty” lanes, Fig.2i). When GST alone was added to LRRK2, no additional bands were noted demonstrating that GST is not phosphorylated by LRRK2. This also validates that LRRK2 retains specificity in this assay, autophosphorylating itself as expected, but not a random protein incubated with it (GST). GST-HuD, was phosphorylated by wild-type LRRK2 and LRRK2 G2019S but not kinase-dead LRRK2 D1994A (Fig.2k). HuB and HuC were phosphorylated by LRRK2 G2019S similarly to HuD (Fig.2h-j, Supplementary Fig.3b).Quantitatively, phosphorylation of HuD was increased 3-fold by LRRK2 G2019S compared to wild-type LRRK2 (Fig.2l), similar to the increased autophosphorylation of LRRK2 observed here, and the increased kinase activity of LRRK2 G2019S kinase on other substrates^10^. Together, this demonstrates that LRRK2 can directly phosphorylate multiple neuronal ELAVs (Fig.2h-j, Supplementary Fig.3b), and their phosphorylation is increased by LRRK2 *in vitro* (Fig.2k,l), in neuronal cell lines (Fig.2e-g) and by *LRRK2 G2019S* in mouse midbrain (Fig.2h-j) using three independent types of experiments (*in vitro* kinase assay, phospho-Threonine antibodies, PhosTag gels).

We sought to identify the sites in HuD phosphorylated by LRRK2. Mass spectrometry was performed on neuronal ELAV immunoprecipitates from mouse brains to identify physiologically relevant phosphorylation sites on HuD for the serine/threonine kinase activity of LRRK2^10^. Phosphorylation at T147, T149 and S223 of neuronal ELAVs was observed across different animals in the mouse brain. To test whether LRRK2 was capable of phosphorylating these sites directly, mass spectrometry was performed on recombinant HuD after incubation with LRRK2 G2019S or kinase-dead LRRK2 with peptide coverage of 65-68%. Phosphorylation of HuD at T149 was consistently increased after incubation with LRRK2 G2019S over three independent replicates (Supplementary Fig.3c). While it is possible that LRRK2 G2019S phosphorylates HuD on additional sites, we focused our attention on T149 because it was the only site both consistently phosphorylated by LRRK2 G2019S *in vitro* and phosphorylated in the mouse brain. T149 is also conserved in HuB, and replaced by a Serine in HuC which could also be phosphorylated by the serine/threonine kinase activity of LRRK2 (Supplementary Fig.3d). T149 and its surrounding sequence is also highly conserved in ELAVs through zebrafish, *C.elegans* and *Drosophila* (Supplementary Fig.3d). T149 is located on an accessible surface of HuD within the RNA-binding RRM2 domain. This suggests T149 is available for phosphorylation by LRRK2 and this could cause the effects on HuD binding to RNA induced by LRRK2 (Fig.1c-l).

To verify that T149 is phosphorylated by LRRK2 G2019S, this site was mutated to alanine. Phosphorylation of HuD T149A by LRRK2 G2019S *in vitro* was strongly reduced compared to wild-type HuD in the *in vitro* kinase assay (Fig.3a). Phosphorylation of HuD T149A was still reduced when a 3-fold excess of the protein was added to the assay (Fig.3a). This demonstrates that *in vitro* LRRK2 G2019S may phosphorylate multiple sites on HuD at a low rate, but the principal site of phosphorylation is T149. In cells, the HuD T149A mutation did not affect levels of HuD, suggesting it has no major impact on protein stability or folding (Fig.3b). Detection of HuD with antibodies specific for phospho-Threonine was also significantly reduced by the T149A mutation (Fig.3c,d). Collectively, this demonstrates that T149 on HuD is phosphorylated in cells, and that this is the dominant site on HuD phosphorylated by LRRK2. Mutation of T149 reduced binding of HuD to target mRNAs including *SNCA*, *BDNF* and *LRRK2* mRNA (Fig.3e), closely resembling the effects of LRRK2 siRNA and kinase inhibitors (Fig.1c-j). This suggests that LRRK2 phosphorylates HuD at T149 to increase its binding to RNA *in vitro*, in cells and mouse midbrain (Fig.1-3).

**Figure 3.**
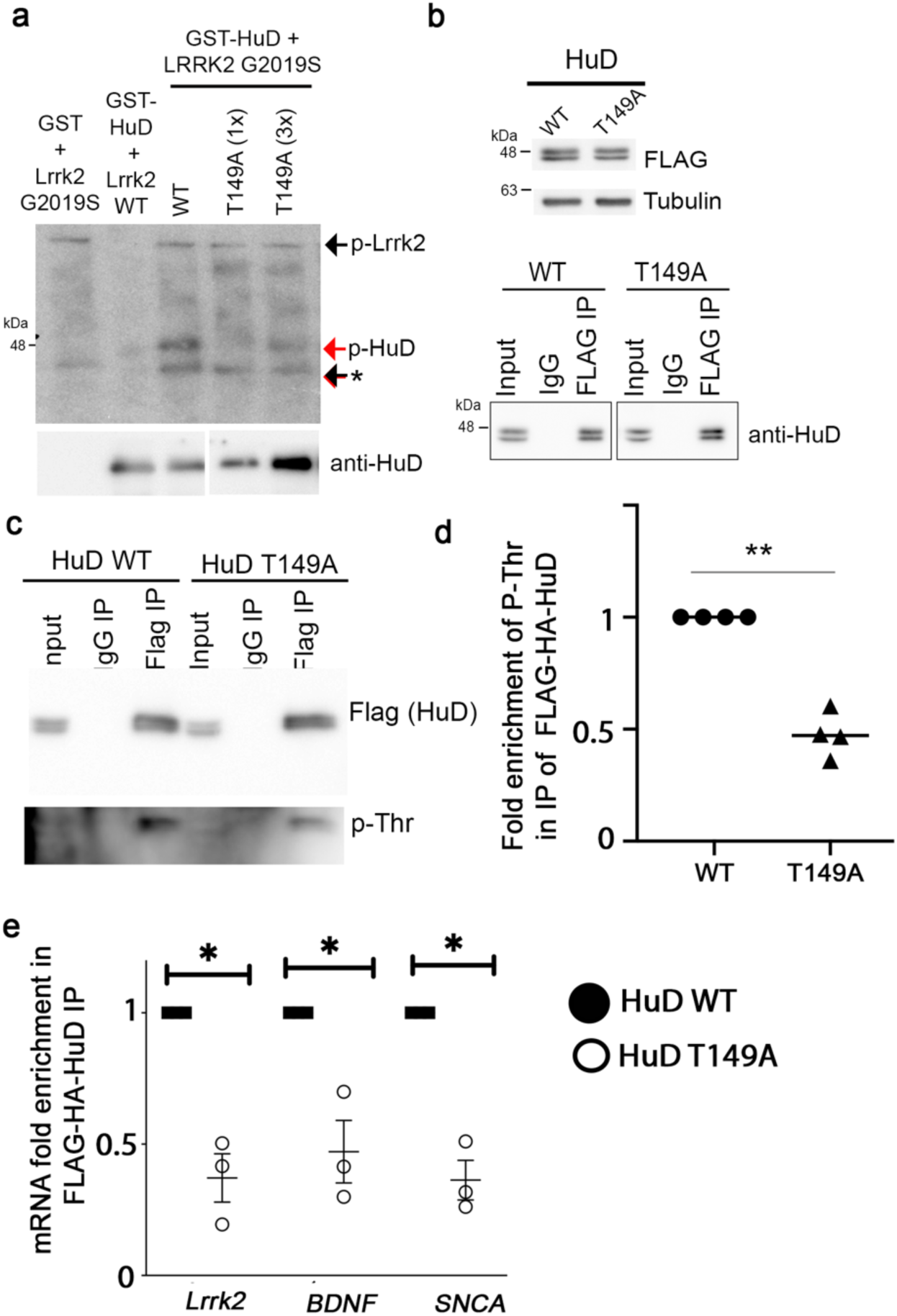
LRRK2 Phosphorylates T149A on HuD to Control its Binding to RNA. **(a)** Autoradiography of HuD phosphorylation with *γ*-32P by LRRK2. SDS-PAGE gel exposed to film after incubation of LRRK2 wild-type or G2019S with *γ*-32P and wild-type HuD, or equal amounts of HuD T149A mutant (1x) or 3-fold as much HuD T149A mutant (3x). Below, Western blot of HuD was run with the identical samples, in parallel, on a separate gel. **(b)** (Above) Western blot of Flag in total cell lysates of cells expressing wild-type or T149A-Flag-HuD, or (Below) immunoprecipates of Flag from the same cells. **(c)** Western blot for phospho-Threonine (p-Thr) in immunoprecipitates of wild-type or T149A Flag-HuD **(d)** Quantification of phosphor-Threonine in immunoprecipitates of wild-type or T149A Flag-HuD over three experiments. One sample t-test **(e)** RT-qPCR quantification of mRNAs in immunoprecipitates of wild-type or T149A Flag-HuD. * p<0.05, one-sample t-test.

### LRRK2 Inhibits Function of HuD in Regulating α-Synuclein and BDNF

Many post-transcriptional regulators, like ELAVs, will impact both mRNA levels and translation of their target mRNAs. We investigated whether LRRK2 by acting on HuD regulates expression of proteins with established relevance to PD: *α*-synuclein and BDNF, since the mRNA encoding these proteins binds to HuD (Fig.1). Expression of HuD or inhibition of LRRK2 alone in neuronal SH-SY5Y cells had no significant impact on protein levels of *α*-synuclein, p21 or BDNF or the mRNAs encoding them (Fig.4a-e). If phosphorylation by LRRK2 inhibits HuD function, then LRRK2 might prevent HuD from exerting effects in SH-SY5Y cells. When LRRK2 was depleted with siRNA, to release HuD from inhibition, levels of BDNF and *α*-synuclein were increased (Fig.6c-e). This shows that HuD promotes production of BDNF and *α*-synuclein but is constitutively inhibited by LRRK2 (Fig.6a-e).

**Figure 4.**
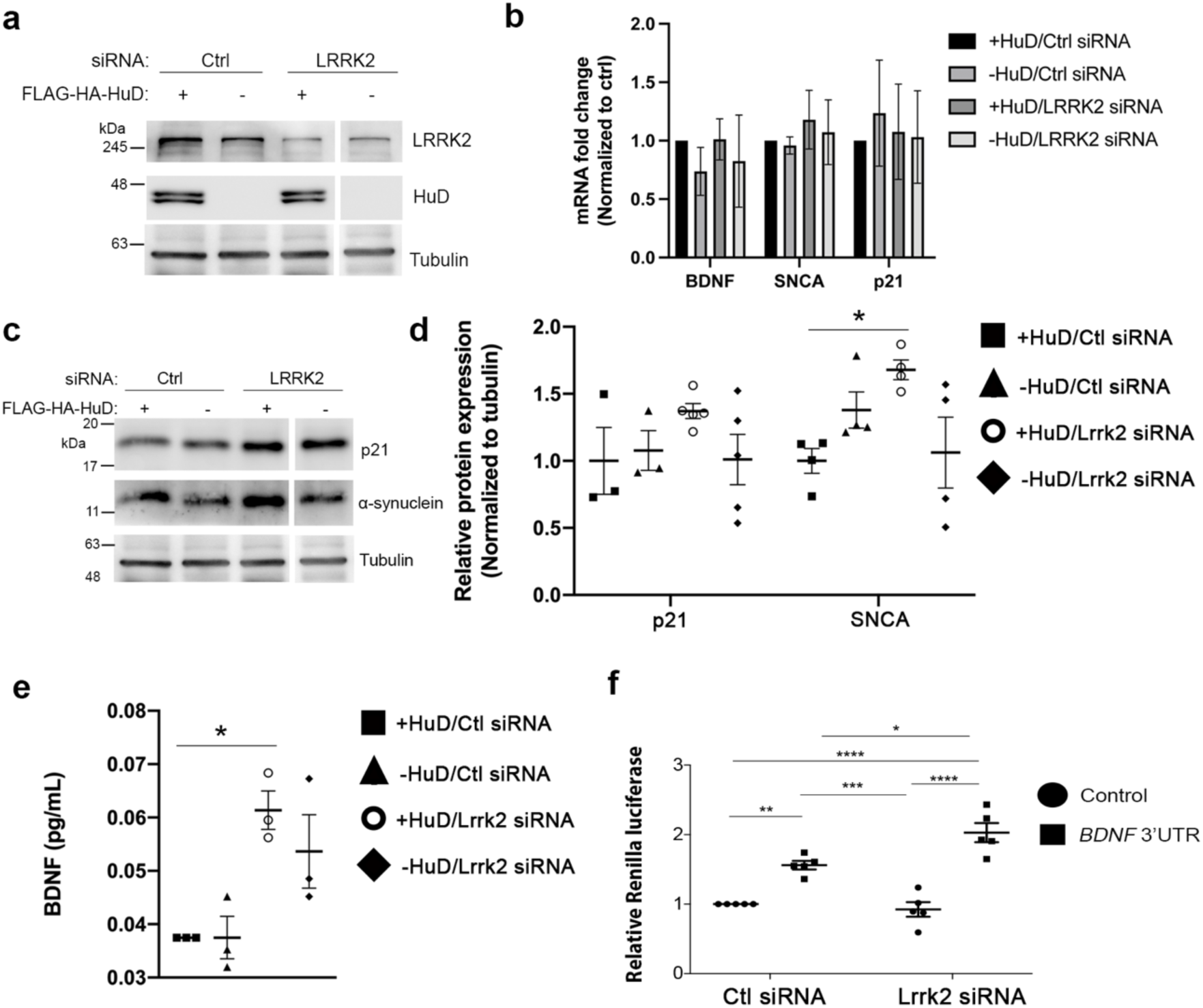
LRRK2 Controls Levels of *α*-Synuclein and BDNF Via Neuronal ELAVs and Their 3’UTR Binding Sites. **(a)** Western blot of levels of LRRK2 and HuD in SH-SY5Y cells after transfection with Flag-HuD or control plasmid, and control siRNA or LRRK2 siRNA **(b)** RT-qPCR for *BDNF, α-synuclein* (SNCA) and *p21* mRNAs in SH-SY5Y cells after transfection with Flag-HuD or control plasmid, and control siRNA or LRRK2 siRNA **(c)** Western blot of levels of *α*-synuclein and BDNF in SH-SY5Y cells after transfection with Flag-HuD or control plasmid, and control siRNA or LRRK2 siRNA. Note that Westerns in (a) and (c) were performed on the same membrane and therefore use the same Tubulin control. **(d)** Quantification of the levels of *α*-synuclein and p21 as in (c). **(e)** ELISA of BDNF in cell supernatants of SH-SY5Y cells after transfection with Flag-HuD or control plasmid, and control siRNA or LRRK2 siRNA **(f)** Luciferase assays of neuronal ELAV binding sites in the *BDNF* 3’UTR in U2OS cells transfected with Flag-HuD and treated with control siRNA or LRRK2 siRNA. The control is an independent section of the *BDNF* 3’UTR that does not contain neuronal ELAV binding sites. * p<0.05, ** p<0.01, *** p<0.001 ANOVA with Tukey’s correction.

To demonstrate that LRRK2 and HuD regulates BDNF through 3’UTR binding sites (Supplementary Fig.2b) luciferase assays were employed. Echoing the effects on protein levels of BDNF, HuD increased translation of luciferase mRNAs containing 3’UTR segments from *BDNF* which harbor known HuD binding sites^40^, and relieving HuD inhibition by depleting LRRK2, further increased production of the BDNF 3’UTR reporter (Fig.6f). These results confirmed that LRRK2 inhibits HuD effects on post-transcriptional gene expression.

If LRRK2 G2019S inhibits neuronal ELAVs, then one might expect LRRK2 G2019S mice to exhibit phenotypes resembling those of a *HuD^-/-^* mouse. Most models of transgenic mice expressing LRRK2 G2019S have very subtle or no pathological phenotype^10, 51^, unlike *HuD^-/-^* mice which exhibit significant motor deficits^48^. We hypothesized that in the absence of a triggering event, the full complement of neuronal ELAVs may be abundant and redundant enough to compensate for their inhibition by LRRK2 G2019S. Others have observed that subtle effects of the loss of *HuD* in mice on mRNA expression and behaviour are amplified when multiple neuronal ELAVs are deleted^33^. Our data, shows that LRRK2 G2019S phosphorylates HuB and HuC like HuD (Supplementary Fig. 3b) regulates their binding to mRNA targets shared with HuD (Fig.1e,i,j, Supplementary Fig.1b-f), and inhibits the effects of neuronal ELAVs on gene expression (Fig.4). We hypothesized that when fewer neuronal ELAVs are present in *HuD^-/-^* mice, that LRRK2 G2019S may inhibit HuB and HuC and lead to more pronounced phenotypes. To test this, *HuD^-/-^* mice were crossed with *LRRK2 G2019S* knock-in mice.

### LRRK2 G2019S Regulates Expression of Neuronal ELAV Targets in the Mouse Midbrain

We first sought to ascertain the impact of LRRK2 G2019S on neuronal ELAVs targets in a genome-wide fashion. Others have reported that neuronal ELAVs (HuB, C and D) can either increase or decrease mRNA stability and translation by binding U-rich sequences in the 3’UTR of mRNAs^33, 52^. To define neuronal ELAV targets, RNA from neuronal ELAV immunoprecipitates (Supplementary Fig.1g) from mouse midbrain was quantified. Neuronal ELAVs bound 2078 RNAs in mouse midbrain with a minimum 4-fold enrichment vs. IgG control including mRNA encoding alpha-synuclein (FDR p>0.01, Supplementary Table 1,2). Neuronal ELAV immunoprecipitates were enriched in binding motifs for neuronal ELAVs and mRNAs previously identified to stringently bind neuronal ELAVs by CLIPseq ^33^ (Fig.5a-b). This validates that the RNA immunoprecipitated with neuronal ELAVs here accurately reproduces previous literature.

**Figure 5.**
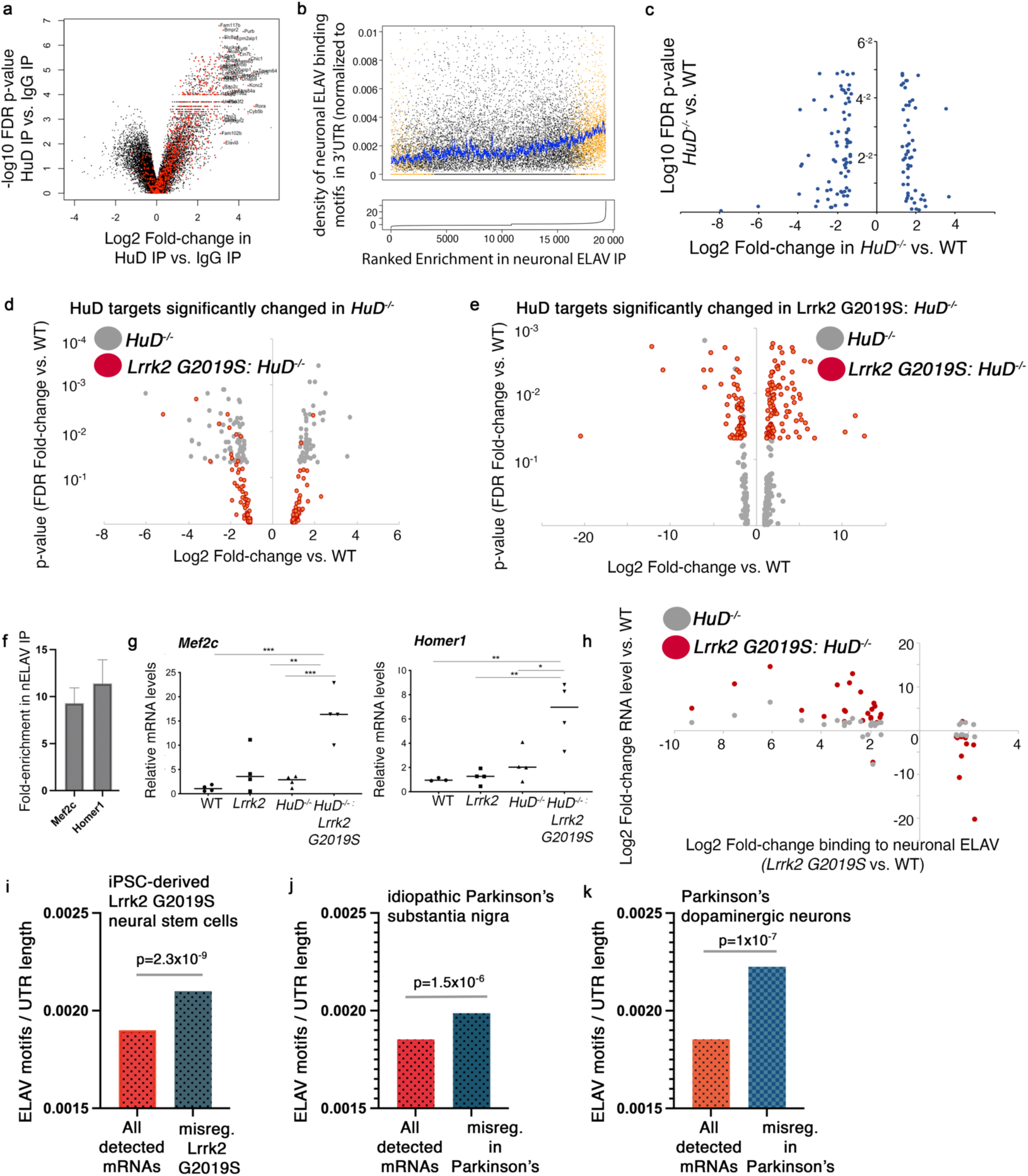
LRRK2 G2019S Exaggerates the Effects of *HuD^-/-^* on Targets of Neuronal ELAVs in the Mouse Midbrain. **(a)** Volcano plot of Log2-enrichment vs. p-value of mRNAs in immunoprecipitates of neuronal ELAVs from mouse midbrain. Red dots, enriched in the present neuronal ELAV immunoprecipitates, represent mRNAs identified as bound to neuronal ELAVs in published neuronal ELAV CLIPseq from cortex^33^ **(b)** Increased density of neuronal ELAV binding motifs (UUU*UUU) ^33^) in mRNAs enriched in neuronal ELAV immunoprecipitates. On the bottom, mRNAs are ordered by their enrichment in neuronal ELAV immunoprecipitates and in the top graph the density of neuronal ELAV binding motifs in these mRNAs are plotted with those significantly enriched or depleted from neuronal ELAV immunoprecipitates colored in orange. The blue line plots the moving average density of neuronal ELAV binding motifs. Note the increase in the blue line when enrichment in neuronal ELAV immunoprecipitates increases (orange dots). **(c)** Inverted volcano plot of fold-change in mRNA levels vs. significance (FDR p-value) in *HuD^-/-^* mice for all mRNAs in neuronal ELAV immunoprecipitates (>2 log2 vs. IgG, FDR p>0.01). Levels of mRNAs bound to neuronal ELAV either increase or decrease in midbrain of *HuD^-/-^* mice. **(d)** Volcano plot of all mRNAs in neuronal ELAV ELAV immunoprecipitates (>2 log2 vs. IgG, FDR p>0.01) that are also significantly altered (p>0.05) in *HuD^-/-^* mice. Changes in neuronal ELAV target levels in *HuD^-/-^* mice is largely lost in *LRRK2 G2019S x HuD^-/-^* mice. **(e)** Volcano plot of all mRNAs in neuronal ELAV ELAV immunoprecipitates (>2 log2 vs. IgG, FDR p>0.01) that are also significantly altered (p>0.05) in *LRRK2 G2019S x HuD^-/-^* mice. Despite being bound to neuronal ELAVs, levels of these RNAs are only altered in *LRRK2 G2019S x HuD^-/-^* mice and rarely in *HuD^-/-^* mice. **(f)** Fold- enrichment of *Mef2c* and *Homer1* mRNAs in IPs of neuronal ELAVs from mouse midbrain normalized to immunoprecipitation with control IgG. **(g)** RT-qPCR quantification of the levels of *Mef2c* and *Homer1* in samples from mouse midbrain of the indicated genotypes. **(h)** Plot of Log2 Fold-change in RNA level vs. Log2- Fold change in binding to neuronal ELAVs (*LRRK2 G2019S* vs wild-type). Only RNAs bound to neuronal ELAVs (>4-fold, FDR>0.01), whose binding to neuronal ELAVs was altered more than Log(2)x1.5 in *LRRK2 G2019S*, and whose expression was significantly altered (p>0.05) in *LRRK2 G2019S x HuD^-/-^* mice are plotted. Note that decreased binding to neuronal ELAVs in *LRRK2 G2109S* results in increased levels of RNA in *LRRK2 G2019S x HuD^-/-^* mice. Conversely, increased binding to neuronal ELAVs in *LRRK2 G2109S* results in decreased levels of RNA in *LRRK2 G2019S x HuD^-/-^* mice. **(i-k)** Density of neuronal ELAV binding motifs (UUUNUUU) in 3’UTRs normalized to 3’UTR length in **(i)** mRNAs misregulated in *Lrrk2 G2019S* iPSC-derived neural stem cells vs. *Lrrk2* corrected wild-type controls^58^ **(j)** among 782 mRNAs misregulated in *substantia nigra* of idiopathic Parkinson’s disease patients^59^ **(k)** among 168 mRNAs misregulated in dopaminergic neurons of Parkinson’s disease patients^60^.

Total RNA from midbrain of wild-type, *HuD^-/-^*, *Lrrk2 G2019S* and *HuD^-/-^* ;*Lrrk2 G2019S* mice was also analyzed (four mice per group, Supplementary Table 1). In mice expressing *Lrrk2 G2019S* no significant changes in RNA abundance were recorded compared to wild-type mice (Supplementary Table 1). In a genome-wide manner *HuD^-/-^* mice and *Lrrk2 G2019S: HuD^-/-^* mice exhibited changes in the levels of many mRNAs (Supplementary Table 1, Supplementary Fig.5). Focusing on targets of neuronal ELAVs, in midbrain from *HuD^-/-^* mice, 127 RNAs that were significantly bound to neuronal ELAVs and were also significantly upregulated or downregulated (FDR >0.05, Figure 5c, Supplementary Table 3). These RNAs were significantly enriched in RNA and DNA binding proteins (Supplementary Table 5), and these may cause effects on the other RNAs that do not bind neuronal ELAVs but are misregulated in *HuD^-/-^* mice (Supplementary Table 1).

LRRK2 can phosphorylate HuB and HuC, like HuD, to regulate their binding to RNAs (Fig.1 e,i,j, 2, Supplementary Fig.1b-f, 3b). This would empower LRRK2 G2019S to either rescue the effect of *HuD^-/-^* on RNA, if it activates HuB and HuC, or alternatively exacerbate the effect of *HuD^-/-^* if it inhibits HuB and HuC. Changes in the levels of the 127 mRNAs significantly bound to neuronal ELAVs and significantly altered in *HuD^-/-^* mice were largely reversed in *Lrrk2 G2019S: HuD^-/-^* mice (Fig.5d, Supplementary Table 3).

A unique subset of 171 RNAs which bound to neuronal ELAVs was not significantly changed in *HuD^-/-^* mice but became significantly different in *Lrrk2 G2019S: HuD^-/-^* mice (Fig.5e, Supplementary Table 4). These RNAs were involved in pathways involved in neurons, synapses and post-synapses (Supplementary Table 6) A model consistent with this, is that because LRRK2 G2019S increases phosphorylation of HuB and HuC (Fig.2h-j, Supplementary Fig.3b) and regulates their binding to RNA (Fig.1) Lrrk2 G2019S can increase HuB and HuC activity and reduce the effects of the loss of *HuD* on one set of RNAs (Fig.5d), while inhibiting the ability of HuB and HuC to regulate another set of targets and exacerbating the effects of *HuD^-/-^* (Fig.5e).

We validated these results. *Mef2c* and *Homer1* mRNAs, bound neuronal ELAVs (Fig.5f). Their levels in total RNA were subtly increased in *HuD^-/-^* mice, and strongly increased in *LRRK2 G2019S: HuD^-/-^* mice as detected in genome-wide analyses (Fig.5g, Supplementary Table 1). *Mef2c* impacts motor function, learning and memory in mice and patients^53, 54^, and *Homer1* regulates survival of dopaminergic neurons in models of PD^55, 56^. This strongly suggests that LRRK2 G2019S alters the subset of RNAs regulated by neuronal ELAVs in *HuD^-/-^* by phosphorylating HuB and HuC *in vitro* and in the mouse midbrain (Fig.2h-j, Supplementary Fig. 3b) to regulate their binding to mRNAs (Fig.1 e,i,j, Supplementary Fig.1b-f). This reinforces data *in vitro* and in cells (Fig.1-4), in a genome-wide manner, that with some transcripts LRRK2 G2019S inhibits the function of neuronal ELAVs, exacerbating effects in *HuD^-/-^* mice, while with other transcripts it activates neuronal ELAVs to minimize the effects in *HuD^-/-^* mice.

To gain insight into the mechanism by which *LRRK2 G2019S* exerts these effects, we examined the effects on RNAs whose binding to neuronal ELAVs was altered in *LRRK2 G2019S* mice vs. wild-type mice. mRNAs whose binding to neuronal ELAVs was decreased by *LRRK2 G2019S* had subtly increased levels in *HuD^-/-^* mice that were further increased by co-expression of *LRRK2 G2019S* (Fig.5h). This suggests that neuronal ELAVs are destabilizing this group of mRNAs and *LRRK2 G2019S* releases them from neuronal ELAVs and this destabilization. In agreement, RNAs whose binding to neuronal ELAVs was increased by *LRRK2 G2019S*, had their levels decreased by *LRRK2 G2019S* in *HuD^-/-^* mice (Fig.5h). Both examples show that *LRRK2 G2019S* changes which RNAs are bound to and destabilized by neuronal ELAVs. Neuronal ELAVs also regulate translation of their targets, and LRRK2 G2019S likely has additional effects on neuronal ELAVs at this level (see above and below).

### mRNAs bound by HuD are misregulated in patients with PD

We queried whether inhibition of neuronal ELAVs may occur in patients with PD. For example, neuronal ELAVs may be inhibited in a broad subset of patients with PD due to over-activation of wild-type LRRK2 as was reported to occur in approximately 90% of patients with sporadic PD^57^ or through other mechanisms. To measure neuronal ELAV inhibition, we queried whether targets of neuronal ELAVs were disproportionately misregulated in patients and patient-derived models of PD compared to controls using public data. 3’UTRs were extracted from all mRNA detected in neural stem cells derived from induced pluripotent stem cells from *LRRK2 G2019S* patients vs. corrected *LRRK2* wild-type^58^. The number of neuronal ELAV binding motifs were normalized to the length of 3’UTRs. Among mRNAs downregulated in *LRRK2 G2019S* neural stem cells there is a significant enrichment in neuronal ELAV binding motifs (UUU*UUU) compared to mRNAs unaffected by *LRRK2 G2019S* (Fig.5i, p=2.3^-9^, proportion test, 0.0021 vs 0.0019 motifs/length). To test whether misregulation of neuronal ELAV binding mRNAs may thus be characteristic of broader groups of idiopathic PD patients we queried this using mRNAs consistently dysregulated in the *substantia nigra* across multiple studies in PD patients^59^. Normalizing to 3’UTR length, the 3’UTRs of 782 unique mRNAs dysregulated in PD patients^59^ were significantly enriched in neuronal ELAV binding motifs compared to all mRNAs detected (Fig.5j, p=1.51x10^-6^, motifs/3’UTR length UUUNUUU: 0.0019873 vs. 0.0018528). To ensure the misregulation of mRNAs containing neuronal ELAV-binding motifs was not due to a change in the cellular composition of degenerating *substantia nigra*, we analyzed data from purified dopaminergic neurons from a broad population of Parkinson’s patients^60^. The 168 misregulated mRNAs in dopaminergic neurons from Parkinson’s patients were also enriched in neuronal ELAV binding motifs compared to all detected mRNAs (Fig.5k, p=1x10^-7^, motifs/3’UTR UUUNUUU: 0.0022255 vs, 0.0018540). Together, this suggests that misregulation of neuronal ELAV targets is characteristic of a broad population of patients with PD both with and without *LRRK2* mutations.

### LRRK2 G2019S Corrects Splicing Defects in HuD^-/-^ *mice*

The results above demonstrate that LRRK2 regulates RNA levels of neuronal ELAV targets in mice and patient cells. Neuronal ELAVs also control mRNA splicing^33^. In mouse midbrain, LRRK2 *G2019S* had no independent effects on splicing of mRNAs that bind neuronal ELAVs (Supplementary Table 7). As previously reported *HuD^-/-^* mice exhibited extensive changes in mRNA splicing, with 1125 mRNAs exhibiting significant differences in splicing (p<0.01, Supplementary Table 8). These mRNAs differentially spliced in *HuD^-/-^* mice were enriched in factors involved in motor function, gait disturbance, brain atrophy and higher mental functions (Supplementary Table 9). Among these alternatively spliced mRNAs, 100 were also bound by neuronal ELAVs and likely directly regulated by HuD (p<0.01, Fig.6a,b, Supplementary Table 10). Remarkably, 92% of these splicing changes of neuronal ELAV targets in *HuD^-/-^* mice were prevented by co-expression of LRRK2 G2019S whether analyzed as p-value or fold-change in splicing (Fig.6a,b, Supplementary Table 10). This effect was validated on a predicted splice variant in *CELF4*. Neuronal ELAVs in the mouse midbrain bound to *CELF4* mRNA (Fig.6c). A long splice isoform of *CELF4* was induced in *HuD^-/-^* mice and this splice variant was reduced by co-expression of *LRRK2* G2019S (Fig.6d). This demonstrates that while phosphorylation by LRRK2 inhibits the function of neuronal ELAVs on mRNA levels and translation (Fig.4-5), it promotes the functions of neuronal ELAVs in mRNA splicing (Fig.6a-d).

**Figure 6.**
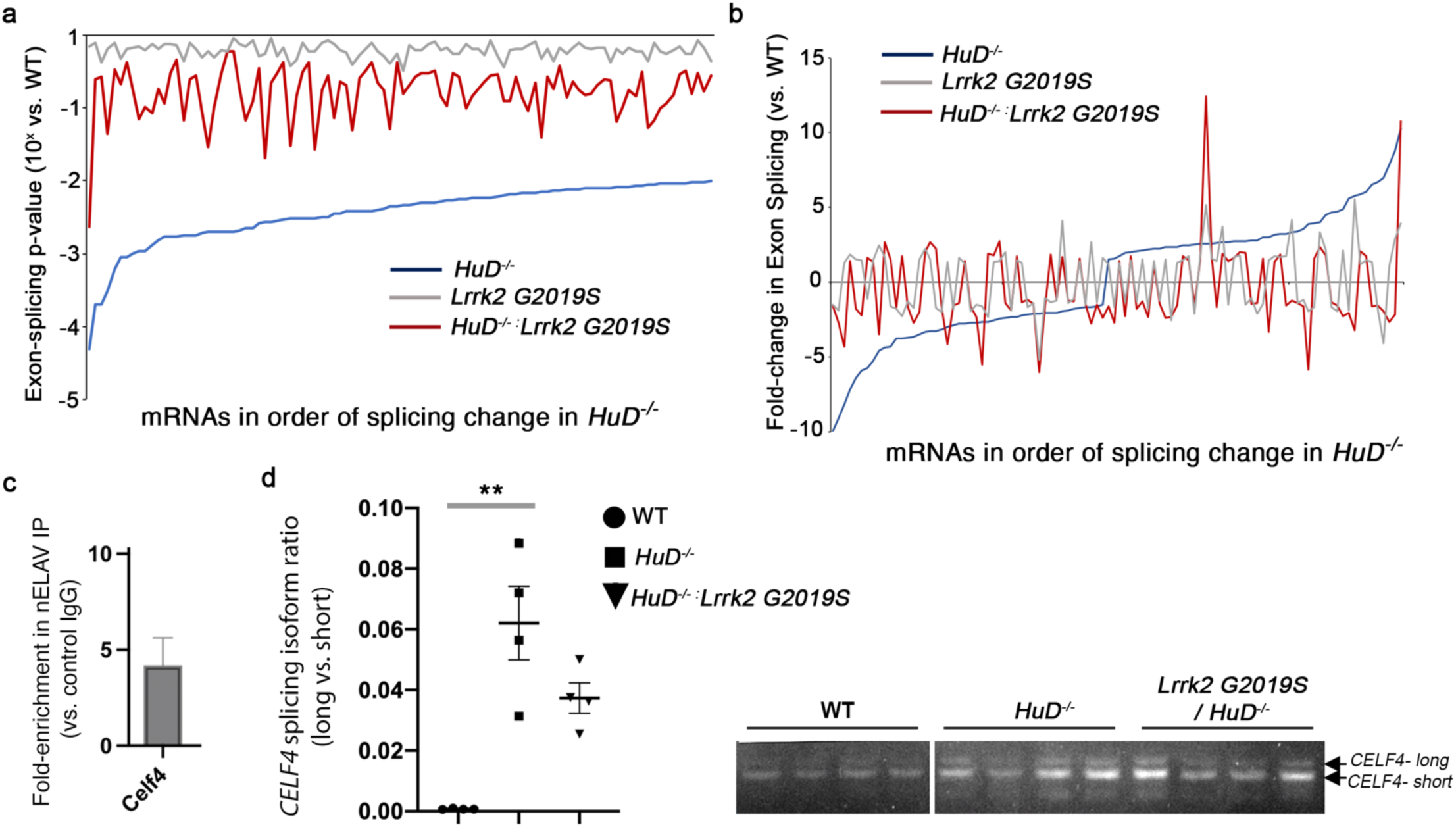
LRRK2 G2019S Minimizes Splicing Defects Caused by Loss of HuD. **(a)** Rank-order of all significant (FDR<0.01) splicing changes in *HuD^-/-^* mouse midbrain by p-values. Note mRNAs whose splicing is significantly altered in *HuD^-/-^* mice are nearly all reversed by *LRRK2 G2019S*. **(b)** Rank-order of all significant (FDR<0.01) splicing changes in *HuD^-/-^* mouse midbrain by fold-change in exon- splicing. Note amplitude of effect on mRNA splicing in *HuD^-/-^* mice is diminished by *LRRK2 G2019S*. **(c)** Quantification of *CELF4* splice variants (long vs. short form) n=4. **(d)** *CELF4* splice variants amplified by RT- PCR and analyzed by agarose gel. Anova with Dunnett’s correction * p>0.01, ** p>0.001

Collectively, these results demonstrate that *Lrrk2 G2019S* may exacerbate some effects in *HuD^-/-^* mice which are due to regulation of mRNA levels, while reducing other effects in *HuD^-/-^* mice which are due to regulation of mRNA levels or splicing by neuronal ELAVs. We evaluated whether neuronal ELAV targets relevant to PD were impacted in a similar manner.. Neuronal ELAVs bind to mRNAs encoding *α*-synuclein and LRRK2 (Fig.1, Supplementary Table 1). Levels of *α*-synuclein were increased in the striatum of *HuD^-/-^* mice and LRRK2 G2019S exacerbated this effect (Fig.7a). In midbrain of *HuD^-/-^* mice, levels of *α*-synuclein were increased and LRRK2 G2019S reduced this effect (Fig.7a). Similarly, BDNF levels increased in *HuD^-/-^* mice and these effects were reduced by LRRK2 G2019S in some brain regions (Fig.7b). Increased LRRK2 abundance and activity are associated with PD. HuD bound mRNAs encoding LRRK2 (Fig.1), suggesting HuD controls expression of LRRK2. Resembling effects on other HuD targets the levels of LRRK2 protein increased in midbrain and striatum of *HuD^-/-^* mice (Fig.7c). Together this demonstrates that neuronal ELAVs regulate expression of multiple proteins with roles in survival of dopaminergic neurons, motor phenotypes and PD (e.g. LRRK2, *α*-synuclein, BDNF, Homer1^55, 56^ and MEF2C^53, 54^) and that LRRK2 G2019S modifies these effects.

**Figure 7.**
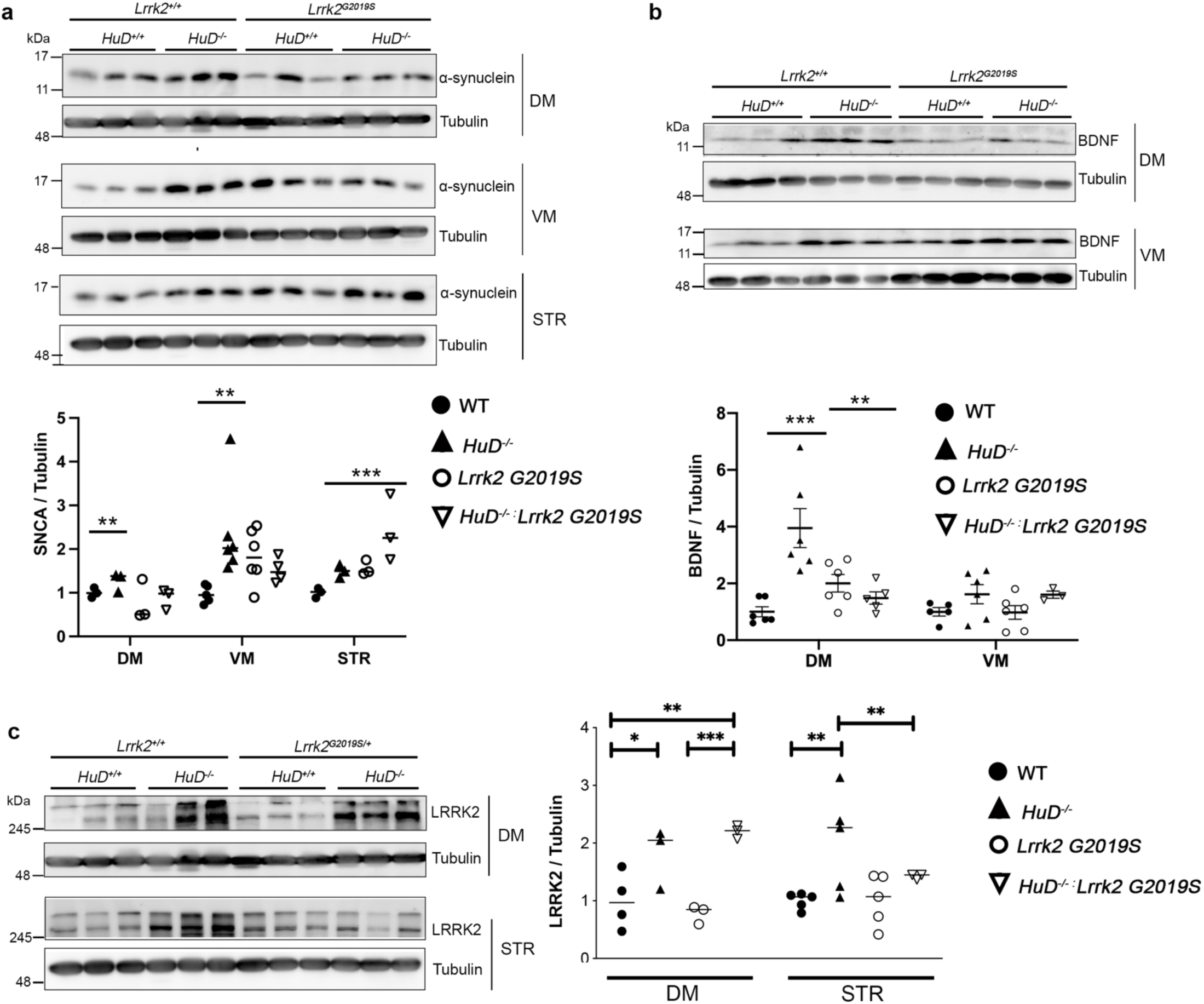
*HuD* and *LRRK2 G2019S* Regulate Alpha-Synuclein, BDNF and LRRK2 Protein Levels in the Mouse Midbrain. **(a,b)** Western blot (top) and quantification (bottom) of *α*-synuclein **(a)** and BDNF **(b)** in wild-type, *HuD^-/-^*, *LRRK2 G2019S,* and *LRRK2 G2019S x HuD^-/-^* mice of 4 weeks of age with dorsal midbrain (DM), ventral midbrain (VM) and striatum (STR) samples normalized to Tubulin (loading control). n=5-6 mice. **(c)** Western blot (left) and quantification (right) of LRRK2 in wild-type, *HuD^-/-^*, *LRRK2 G2019S,* and *LRRK2 G2019S x HuD^-/-^* mice of 4 weeks of age with dorsal midbrain (DM), and striatum (STR) samples normalized to Tubulin (loading control). n=5-6 mice. Note that blots for Lrrk2 and alpha-synuclein were performed on the same membranes and therefore use the same Tubulin control. * p<0.05, ** p<0.01, *** p<0.001 ANOVA with Tukey’s correction where multiple groups were compared, t-test (two tailed assuming unequal variance) where one group was compared.

A key phenotype of PD is loss of dopaminergic neurons. Stereological counting of Tyrosine Hydroxylase+ dopaminergic neurons in serial sections of the substantia nigra of mice at 12 months showed insignificant differences in mice with a *LRRK2 G2019S HuD^-/-^* mice alone (Fig.8a,b). However, *HuD^-/-^*;*Lrrk2 G2019S* mice showed a modest but significant reduction in dopaminergic neurons in the substantia nigra in line with aggravated dysregulation of some neuronal ELAV targets in these mice (Fig.4,5,7a-c). Onset of motor symptoms in PD patients and mouse models usually occurs after loss of 30-50% of dopaminergic neurons in the substantia nigra^61, 62^, similar to the amount of loss of these neurons observed in *HuD^-/-^*;*Lrrk2 G2019S* mice. Another key phenotype of PD is motor deficits such as changes in gait. *HuD^-/-^* mice appeared to have altered gait, but this only reached significance in *HuD^-/-^*;*Lrrk2 G2019S* mice (Fig.8a). Similarly, overall movement assessed by beam breaks tended to decrease in *HuD^-/-^* mice, but this reached significance in *HuD^-/-^*;*Lrrk2 G2019S* mice (Fig.8b). This suggests that loss of *HuD* sensitizes mice to the appearance of PD phenotypes including loss of dopaminergic neurons and motor deficits that can be triggered by LRRK2 G2019S.

**Figure 8.**
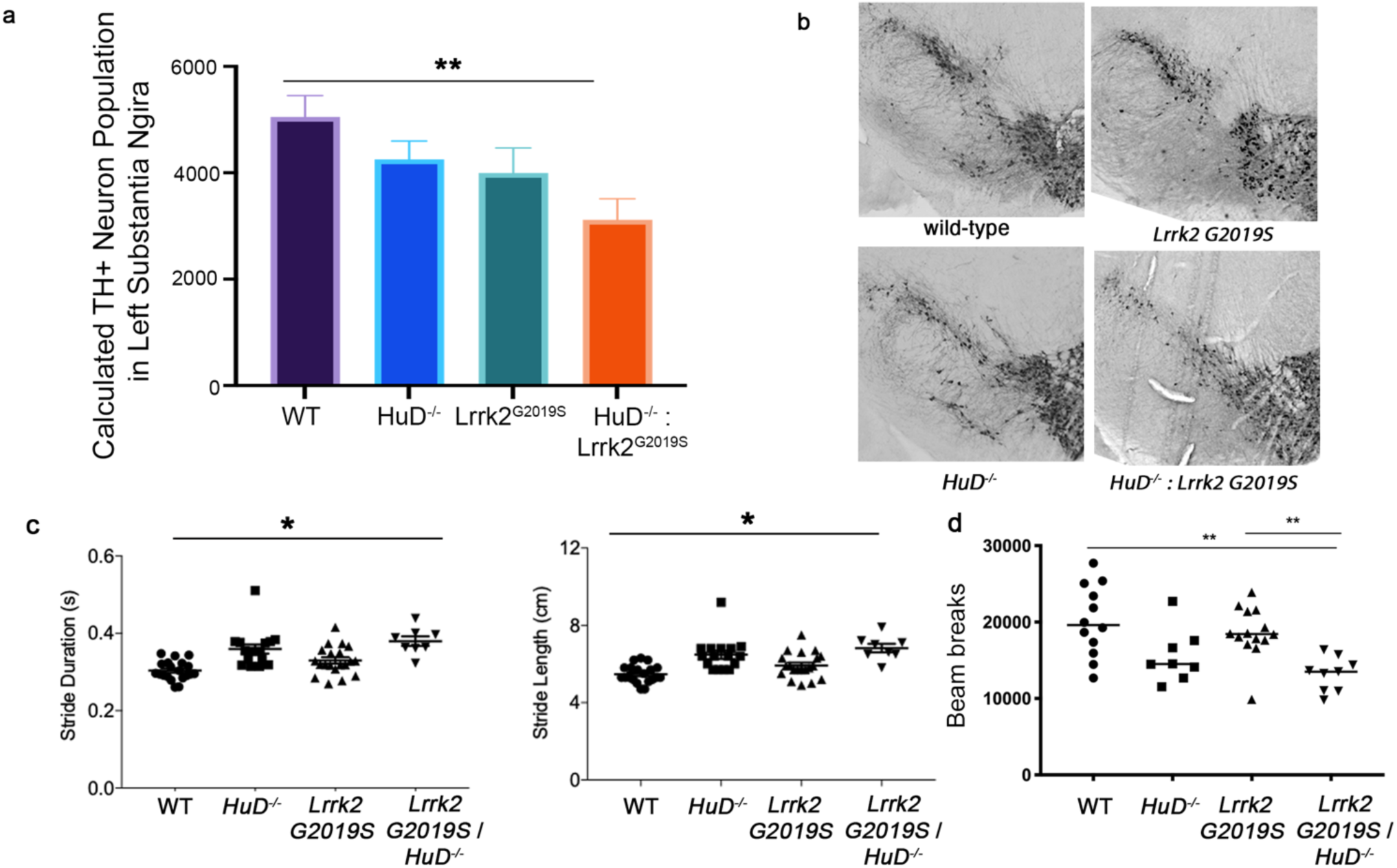
*LRRK2 G2019S x HuD^-/-^* mice Exhibit Loss of Dopaminergic Neurons and Motor Deficits. **(a,b)** Stereological counting of Tyrosine-Hydroxylase+ dopaminergic neurons in serial sections of one half of the substantia nigra. **(a)** Quantification of representative images **(b)** of dopaminergic neurons in one side of the substantia nigra of wild-type, *HuD^-/-^*, *LRRK2 G2019S*, *LRRK2 G2019S* x *HuD^-/-^* mice. n=6 mice per genotype. * p<0.05, Dunnett’s post-hoc **(a)** Quantification of stride duration and length in wild-type, *HuD^-/-^*, *LRRK2 G2019S*, *LRRK2 G2019S* x *HuD^-/-^* mice at 10 weeks of age using a DigiGait apparatus. n>8 mice per genotype. **(b)** Quantification of total horizontal and vertical movement of wild-type, *HuD^-/-^*, *LRRK2 G2019S*, *LRRK2 G2019S* x *HuD^-/-^* mice at 26 weeks of age. * p<0.001, ANOVA with Tukey’s correction.

Here, we demonstrate that neuronal ELAVs including HuB, HuC and HuD are phosphorylated by LRRK2, and that this regulates their binding to mRNA. LRRK2 inhibits their activity in control of mRNA and protein abundance in cells, mouse models and patient samples. This results in misregulation of many mRNA targets, *α*-synuclein and *Lrrk2* in *HuD^-/-^* mice that can be exacerbated by LRRK2 G2019S. LRRK2 G2019S, possibly by potentiating the inhibition of neuronal ELAVs in *HuD^-/-^* mice induces loss of dopaminergic neurons and motor deficits associated with PD.

## DISCUSSION

Our data demonstrate that LRRK2 can directly phosphorylate neuronal ELAVs *in vitro,* and LRRK2 kinase activity increases neuronal ELAV phosphorylation using recombinant proteins, neuronal cell lines and the mouse midbrain as assessed with phospho-threonine antibodies, radioactive kinase assays, and PhosTag gels. This demonstrates that neuronal ELAVs are a direct and physiological target of LRRK2 in tissues relevant to PD.

Like many proteins which post-transcriptionally regulate gene expression, ELAV family proteins regulate both mRNA stability and translation through processes that intimately link translational inhibition and mRNA decay. In these cases, effects on mRNA levels can often be more subtle than the effects on protein levels. This is likely why HuD associates with hundreds, or even thousands of mRNAs, but the mRNA levels of only about 10% of these are measurably changed in *HuD^-/-^* mice (Supplementary Table 1) or in cells (Fig.4) but differences are observed in protein levels of HuD targets like alpha-synuclein, BDNF and LRRK2 itself (Fig.4,7).

LRRK2 phosphorylates HuD on T149, a major site of HuD phosphorylation in the mouse brain defined by mass spectrometry. T149 is on an exposed loop in its RRM2 domain which mediates binding to U-rich RNAs^36, 63^. Stable binding of HuD to U-rich RNAs requires all three RRMs, and notably RRM2^63^, providing a mechanism by which phosphorylation of HuD on T149 in RRM2 can regulate its binding to mRNA. HuD has been documented to promote mRNA stability and destabilization^33, 37, 42, 50, 64, 65^, and to promote translation through binding to eIF4a^35^ or Y RNAs^52, 66^. The RRM2 and its neighboring linker region to RRM3 also regulate its effects on translation^35, 52, 66^. The RRM2 domain thus appears to be central to most of the effector functions of HuD, and a potent point for regulation by a kinase like LRRK2. While T149 appears to be the principal site of phosphorylation for LRRK2 on HuD, it is possible that phosphorylation of HuD on additional sites contributes to the observed effects in cells and mice.

In agreement with previous publications^33, 52^, we found that mRNAs directly bound and regulated by neuronal ELAVs were divided into two pools which either decreased or increased in the mouse midbrain of *HuD^-/-^* mice (Fig.5c). Notably LRRK2 G2019S amplified the effects of *HuD^-/-^* on levels of some mRNAs while rescuing the effect of *HuD^-/-^* on the levels or splicing of another set of mRNAs (Fig.5d,e, 6). These effects were paralleled in impacts on levels of *α*-synuclein, BDNF and LRRK2 in mouse brain (Fig.7). In some brain regions LRRK2 G2019S exacerbated the effects on levels of these HuD targets in *HuD^-/-^* mice, while in other brain regions *LRRK2 G2019S* reversed the effects on protein levels of HuD targets in *HuD^-/-^* mice.

Changes in the levels of targets of neuronal ELAVs at the level of RNA and protein were accompanied by changes in the binding of the mRNAs to neuronal ELAVs controlled by LRRK2 kinase activity in both cells and the mouse midbrain (Fig.1-4, 5h). A model consistent with our data is that LRRK2 phosphorylates neuronal ELAVs to alter the subset of mRNAs which they bind and regulate. LRRK2 kinase activity phosphorylates HuB, HuC and HuD both *in vitro* and in the mouse brain (Fig.2h-j, Supplementary Fig.3b), regulates binding of each of them to mRNA targets (Fig.1 c-j, Supplementary Fig.1b-f) and regulates the ability of neuronal ELAVs to control expression of its target mRNAs (Fig.4,5).

The hyperactive LRRK2 G2019S hyper-phosphorylates neuronal ELAVs (Fig.2) rescuing the effects of *HuD^-/-^* on some mRNAs while exacerbating its effects on others. One might expect that *Lrrk2 G2019S* mice would exhibit at least in part the motor phenotypes of *HuD^-/-^* mice. We observed control of neuronal ELAV function by LRRK2 in contexts where expression of neuronal ELAVs was limiting. For example, experiments in cell culture were performed in the neuroblastoma cell line SH-SY5Y where endogenous neuronal ELAV expression was low. In the brain, three neuronal ELAVs are expressed, frequently at high levels, while LRRK2 G2019S levels in neurons are low. In this context, the effects of LRRK2 G2019S on neuronal ELAVs is probably often compensated for by the abundance of neuronal ELAVs. When levels of neuronal ELAVs were reduced in *HuD^-/-^* mice, LRRK2 G2019S increased the phosphorylation of the remaining neuronal ELAVs (Supplementary Fig.3b, Fig.2h-j) further inhibiting neuronal ELAV activity on mRNA expression (Fig.4) and levels of BDNF and *α*-synuclein in some brain regions (Fig.7). This suggests that in patients LRRK2 hyperactivity will preferentially elicit neuronal ELAV-dependent phenotypes in regions where neuronal ELAVs are less abundant. Intriguingly, studies in mouse and human have noted that neuronal ELAVs are expressed at low levels in the midbrain^67–69^, the region principally affected in PD, and tend to decrease further with age^69^.

The capacity of neuronal ELAVs to bind and regulate mRNAs encoding LRRK2 and *α*-synuclein (Fig.1,7, Supplementary Table 1) could control levels of these proteins which are critical for PD. LRRK2 kinase activity has previously been shown to regulate the levels of LRRK2 protein^70^ and *α*-synuclein pathology in mice^73, 7172^. This suggests that LRRK2 G2019S could contribute to the accumulation of *α*-synuclein, the acceleration of *α*- synuclein pathology, and LRRK2-dependent pathology by inhibiting neuronal ELAVs.

Mouse models expressing PD-linked mutations in LRRK2 at near physiological levels, such as the mice studied here, have shown minimal or no loss of dopaminergic neurons or motor phenotypes^70, 74^. The combination of *Lrrk2 G2019S* and *HuD^-/-^* induced PD-relevant phenotypes including loss of dopaminergic neurons, accumulation of striatal *α*-synuclein in some tissues, and motor phenotypes (Fig.7,8). LRRK2 G2019S effects on proteins other than neuronal ELAVs likely contribute to these phenotypes. However, our data demonstrates that LRRK2 G2019S increased phosphorylation of neuronal ELAVs in the mouse brain (Fig.2e-g) and specifically exacerbated changes in expression of one pool of mRNAs bound by neuronal ELAVs in *HuD^-/-^* cells while rescuing the effects of *HuD-/-* on another pool of their targets. Neuronal ELAV targets were also selectively misregulated in sporadic PD and neurons derived from patients carrying LRRK2 G2019S mutations. Together, this suggests that at least part of the PD-relevant phenotypes induced by LRRK2 G2019S in *HuD^-/-^* mice are due to inhibition of other neuronal ELAVs.

Several molecular mechanisms have been proposed to be a root cause of PD including disruptions in mitochondrial dynamics, vesicle-trafficking, autophagy, inflammation and accumulation of *α*-synuclein. Recent literature has demonstrated that LRRK2 can phosphorylate Rab proteins involved in organelle trafficking to impact some of these pathways in cell models^23^. Neuronal ELAV proteins by binding and regulating hundreds of mRNAs also regulate mitochondrial dynamics^75^, autophagy^76^, inflammation^77^ and *α*-synuclein and LRRK2 expression (Fig.7). Indeed, we also identified several Rab proteins, including Rab10 among neuronal ELAV targets (Supplementary Table 1). Alongside its regulation of Rabs, LRRK2 could also impact this broad range of pathways with roles in PD in part by phosphorylating ELAV proteins to control their mRNA targets. The association of single-nucleotide polymorphisms in the *HuD* locus to PD further supports the hypothesis that alterations in neuronal ELAV expression or function could contribute to PD^29–31^. This raises the possibility that along with Rab proteins, inhibition of neuronal ELAV function could be a consequential target of LRRK2 kinase, via which it causes PD. If LRRK2 kinase activity is increased in a large majority of patients with idiopathic PD as reported^57^, this mechanism could contribute to PD in many patients.

Are neuronal ELAVs actually inhibited in patients with PD? We demonstrated that neuronal ELAV target mRNAs were selectively dysregulated across studies of iPSC-derived neural stem cells from patients, *substantia nigra*, and isolated dopaminergic neurons from PD patients. This provides evidence that neuronal ELAVs are inhibited in both patients with *LRRK2 G2019S* mutations and the large majority of patients with idiopathic PD, independent of any change in cell composition of tissues. Together, this suggests that loss of neuronal ELAV function, by LRRK2 phosphorylation or other mechanisms, may be common in PD and contribute to its characteristic pathology.

## Acknowledgements

ON was funded by the Toth Family Fellowship (2017) and the Audrey Grant Scholarship (2018) from the Parkinson’s Research Consortium (Ottawa). AP was funded by NSERC Graduate Student Scholarship. Research in the Gibbings and Park lab was funded by an award from the Canadian Institutes for Health Research. The authors thank Mirela Barclay and Christine Luckhart of the University of Ottawa Animal Behaviour Core Facility for performing behavioural testing on mice at 26 weeks of age. The authors thank Novartis for the kind gift of LRRK2 G2019S mice.

## Author Contributions

AP performed and analyzed all work in cell lines unless otherwise noted, immunoprecipitations and western blots from mice, mutagenesis of HuD and immunoprecipitation of mutant HuD. ON performed and analyzed luciferase assays, PhosTag gels, phosphorylation assays on HuB, HuC and mutant HuD, stereological counting of TH+ neurons, RT-PCR for splicing and microarray validation. ARC performed gel shift assays and designed splicing primers. MTT and AS helped generate mice and performed dissections. GP processed microarray data and performed analysis of neuronal ELAV motifs in microarray data and public datasets. AS performed RT-qPCR validation of microarray data. HG performed proximity ligation assays, immunoprecipitation of LRRK2 and HuD, and HuD phospho-threonine. JT helped generate HuD mutants. HO generated and provided HuD knockout mice. LTM helped analyze mass spectrometry data. JFC analyzed phosphorylation sites on neuronal ELAVs. BJ read and commented on the manuscript. DSP and PM conceived the project and helped design and interpret certain experiments. DG conceived the project, designed and interpreted experiments including analysis of microarrays and patient data and wrote the manuscript.

## Supplementary Figures

**Supplementary Figure 1.**
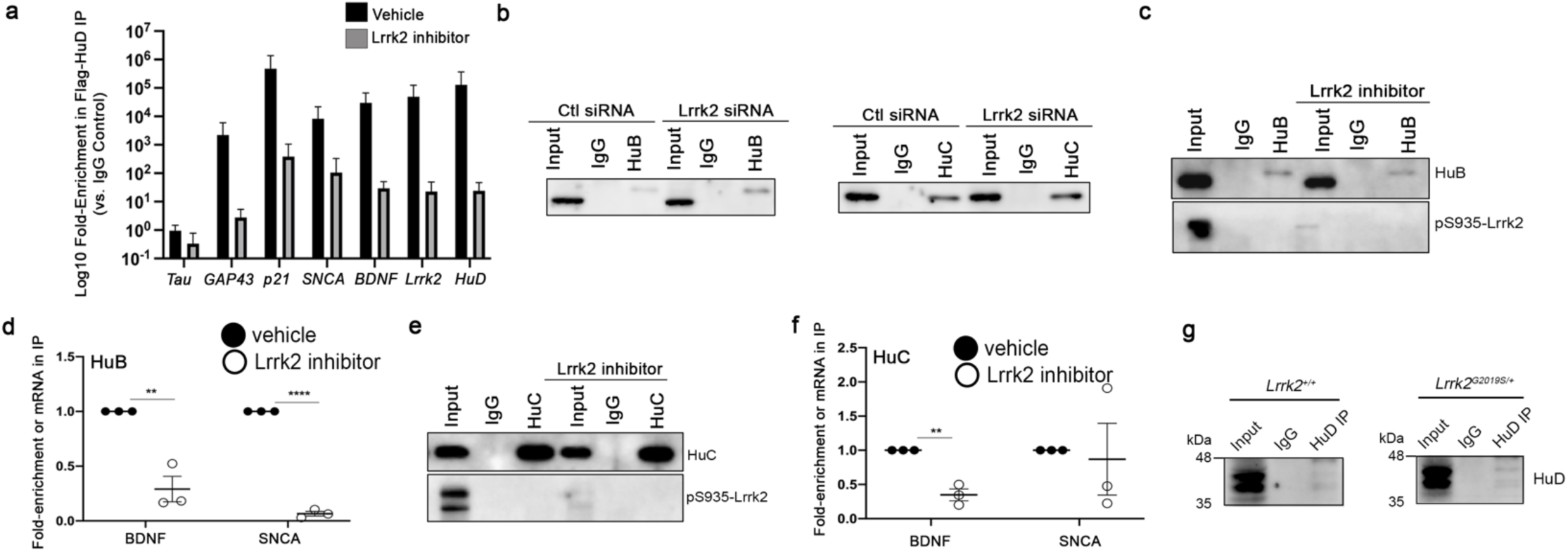
(a) Log10 fold-enrichment of mRNAs in Flag-HuD immunoprecipitated (vs. IgG control immunoprecipitate) after treatment of cells with vehicle of LRRK2 kinase inhibitor. **(b)** Western blot of Flag in immunoprecipitates of Flag-HuB or Flag-HuC from SH-SY5Y cells treated with control siRNA or LRRK2 siRNA. **(c-f)** Effect of LRRK2 kinase inhibitors on mRNA binding to Flag-HuB **(c,d)** and Flag-HuC **(e,f)**. **(c,e)** Western blot of Flag and LRRK2 phosphoserine-935 in SH-SY5Y cell lysates and immunoprecipitates of Flag from cells transfected with Flag-HuB **(c)** or Flag-HuC **(e)**. **(d,f)** Quantification of RNAs by RT-qPCR in immunoprecipitates of Flag-HuB **(d)** or Flag-HuC **(f)** from SH-SY5Y cells treated with vehicle or MLi-2 LRRK2 kinase inhibitor over 2-3 independent experiments. * p<0.05, ** p<0.01, *** p<0.001 ANOVA with Tukey’s correction. **(g)** Western blot of neuronal ELAVs in immunoprecipitates of neuronal ELAV monoclonal antibody or isotype control antibody from wild-type and *LRRK2 G2019S* mouse midbrain.

**Supplementary Fig. 2.**
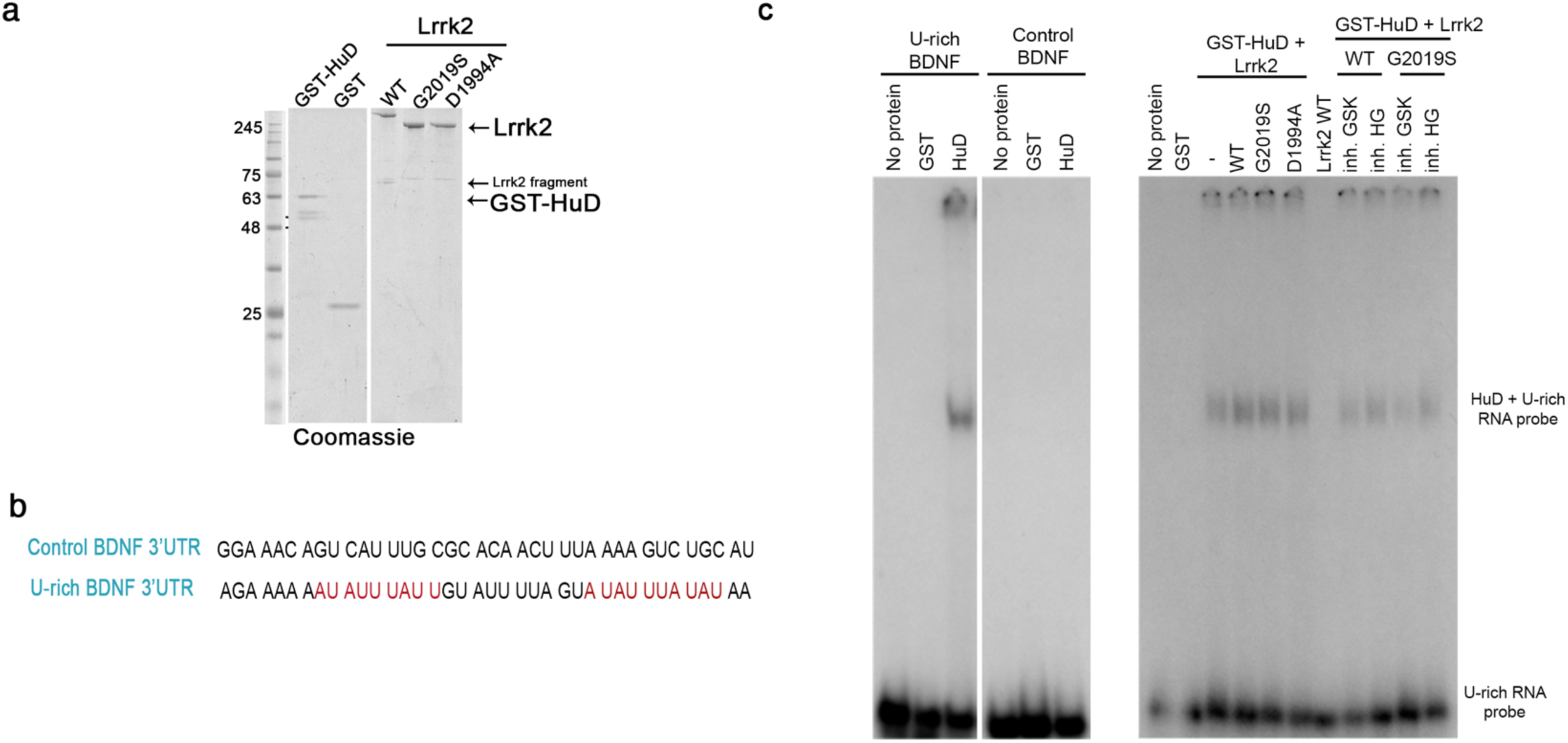
Gel-shift Assays Show LRRK2 Kinase Activity Promotes Binding of HuD to RNA. **(a)** Coomassie total protein stain of recombinant GST, GST-HuD, LRRK2 WT, LRRK2 G2019S and LRRK2 D1994A. **(b)** Model of synthetic RNAs used in experiments Figure 2j and Supplementary Figure 1d. RNA derived from a neuronal ELAV binding segment of the *BDNF* 3’UTR which contains U-rich motifs or an independent segment of the *BDNF* 3’UTR that is not regulated by neuronal ELAVs ^47^. **(c)** Representative autoradiographic images of gel-shift assay using RNA probes as in (b) labeled with *γ*-32P and incubated with GST, GST-HuD, and LRRK2 variants +/- LRRK2 kinase inhibitors.

**Supplementary Figure 3.**
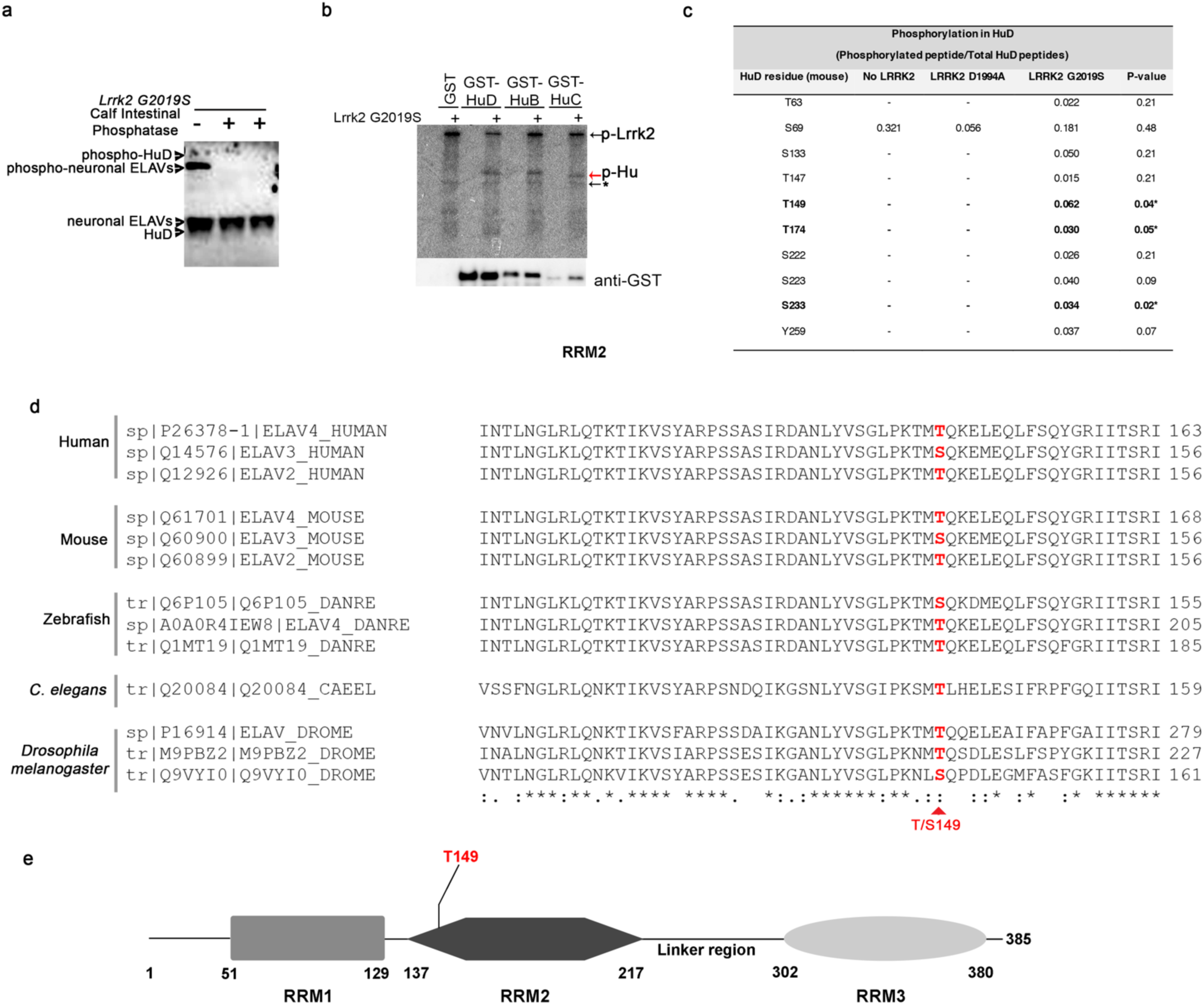
Phosphorylation of neuronal ELAVs by LRRK2. **(a)** Western blot of neuronal ELAVs from the midbrain of *LRRK2 G2019S* mice separated on a PhosTag gel. Equal amounts of samples were left untreated or treated with Calf Intestinal Phosphatase to dephosphorylate proteins and validate detection of phosphorylated neuronal ELAVs on PhosTag gels. **(b)** Autoradiographic exposure of SDS-PAGE gel after incubation of recombinant GST-HuB, -HuC, and -HuD with vehicle or LRRK2 G2019S in the presence of γ-32P. Arrows highlight autophosphorylated LRRK2 or (*) non- specific bands (potentially autophosphorylated fragments of LRRK2 also observed by Coomassie total protein stain (Supplementary Fig. 2a). **(c)** Quantification of recombinant HuD phosphorylation on specific sites by MS/MS (phosphorylated peptide count/Total HuD peptide count) after incubation alone, with kinase-dead LRRK2 D1994A or with LRRK2 G2019S. Peptides significantly phosphorylated after incubation with LRRK2 G2019S vs. D1994A are shown in bold. **(d)** Alignment of mouse, human, zebrafish, *C.elegans* and *Drosophila ELAV* homologues highlighting conservation of LRRK2 phosphorylation site at T149. **(e)** Domain map of neuronal ELAVs indicating the location of T149.

**Supplementary Figure 4.**
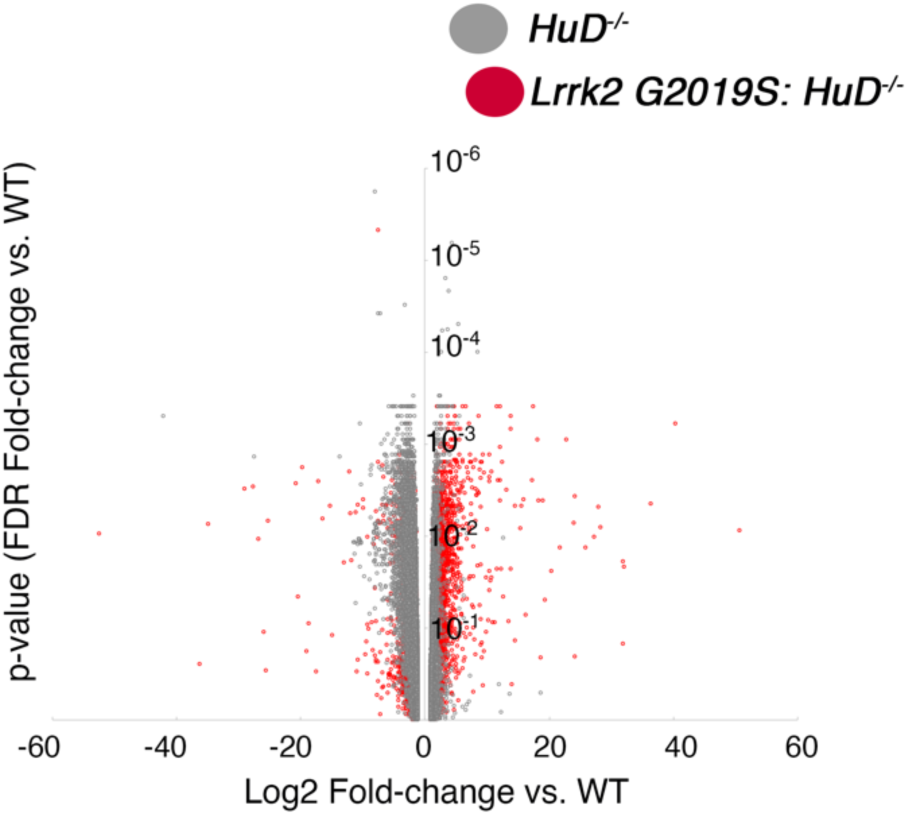
mRNA expression in midbrain of *Lrrk2 G2019S:HuD^-/-^* and *HuD^-/-^* mice. Volcano plot of the Log2 fold-change in mRNA levels (x-axis) vs. the p-value of the fold-change in *Lrrk2 G2019S:HuD^-/-^* and *HuD^-/-^* mice.

## Supplementary Dataset 1

**Predicted binding sites for neuronal ELAVs in the 3’UTR of human and mouse *Lrrk2* and *Snca* mRNAs.** Neuronal ELAVs binding sites have been defined by CLIPseq^33^ as UUUNUUU, where N is any base. Previously, neuronal ELAVs have been defined to bind AU-rich sequences. Here, neuronal ELAV binding sites defined by CLIPseq are in red text in bold and underlined. Other putative binding sites are in red text.

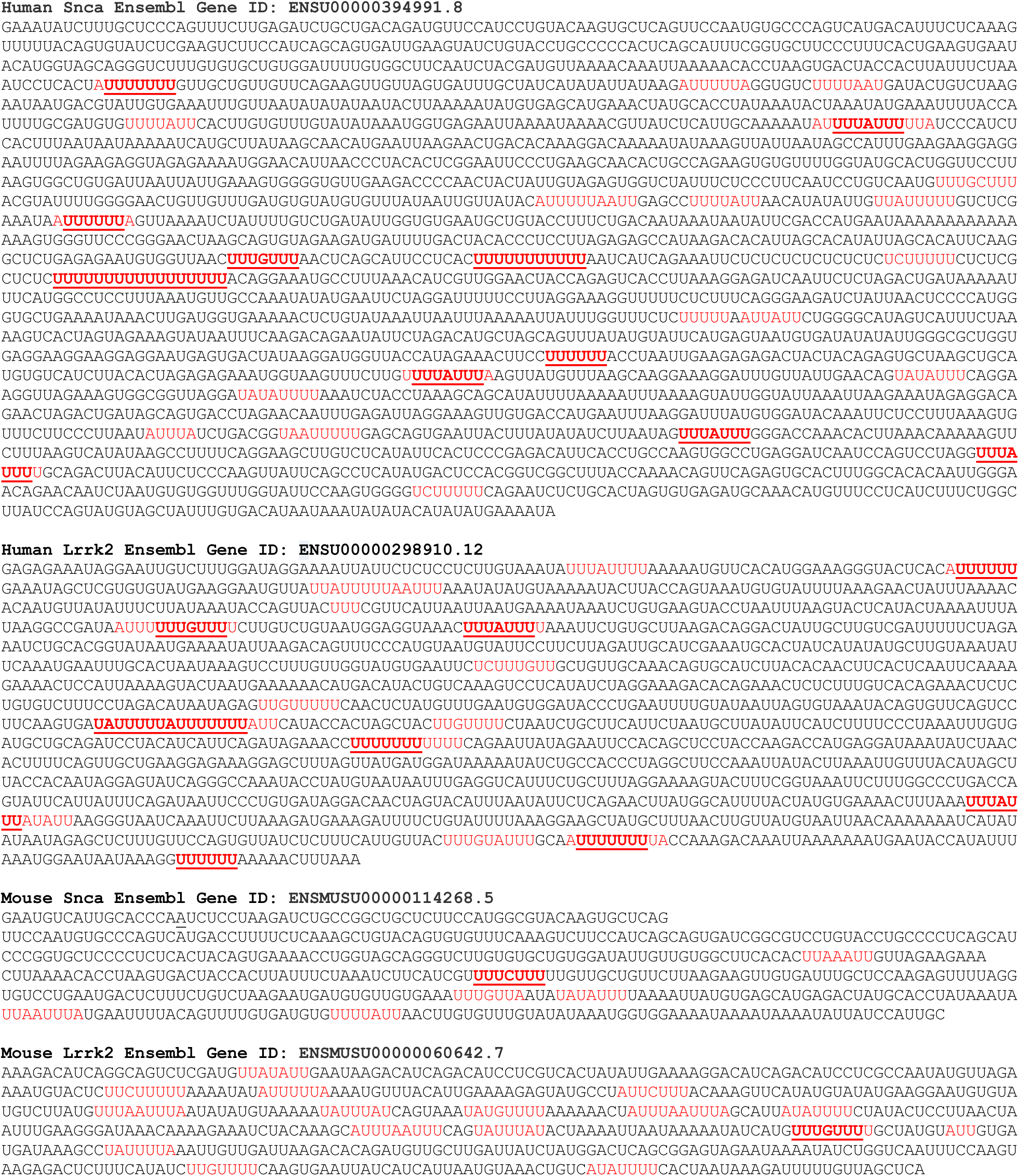

### Supplementary Tables

Supplementary **Tables 1-6.** Impact of HuD and LRRK2 G2019S on RNA Levels

**Supplementary Table 1**. All RNA Expression Data.

**Supplementary Table 2**. All RNAs Bound to Neuronal ELAVs (>log2, FDR<0.01).

**Supplementary Table 3**. All RNAs Bound to Neuronal ELAVs (>log2, FDR<0.01) and Differentially Regulated in *HuD^-/-^* vs. WT midbrain.

**Supplementary Table 4**. All RNAs Bound to Neuronal ELAVs (>log2, FDR<0.01) and Differentially Regulated in *LRRK2 G2019S x HuD^-/-^* vs. WT midbrain (FDR<0.05).

**Supplementary Table 5**. GO-Term analysis of All RNAs Bound to Neuronal ELAVs (>log2, FDR<0.01) and Differentially Regulated in *HuD^-/-^* vs. WT midbrain (FDR<0.05).

**Supplementary Table 6**. GO-Term analysis of All RNAs Bound to Neuronal ELAVs (>log2, FDR<0.01) and Differentially Regulated in *LRRK2 G2019S x HuD^-/-^* vs. WT midbrain (FDR<0.05).

Supplementary **Tables 7-10.** Impact of HuD and LRRK2 G2019S on RNA Splicing

**Supplementary Table 7**. All Splicing Data including exon identifications

**Supplementary Table 8**. All RNAs Differentially Spliced in *HuD^-/-^* vs. WT midbrain (FDR<0.01).

**Supplementary Table 9**. GO-Term analysis of All RNAs Differentially Spliced in *HuD^-/-^* vs. WT midbrain (FDR<0.01).

**Supplementary Table 10**. All RNAs Bound to Neuronal ELAVs (>log2, FDR<0.01) and Differentially Spliced in *HuD^-/-^* vs. WT midbrain (FDR<0.01).

## Materials and Methods

### Cell culture, lysis, transfection and treatments

Mouse neuroblastoma Neuro2A cells and human neuroblastoma SH-SY5Y cells were cultured in DMEM (Wisent) supplemented with 10% fetal bovine serum (FBS), 100U/mL penicillin and 100ug/mL streptomycin at 37°C with 5% CO2. Transient DNA transfection was performed 24 hours post-seeding in low-serum (2-5% FBS), antibiotic-free DMEM using Lipofectamine 2000 (Catalogue 11668-019, Thermo Fisher Scientific). FLAG/HA-HuD was expressed using the pFRT-DestFLAGHA_HuD plasmid (Addgene #65760); a gift from Thomas Tuschl (Rockefeller University, NY, USA), comparing to a control transfection with pcDNA3-EGFP plasmid (Addgene #13031) a gift from Doug Golenbock. Silencer® Select siRNAs (Thermo Fisher Scientific; Appendix - Table 1) were transfected at a concentration of 10nM using RNAiMax (Catalogue 13778150, Thermo Fisher Scientific). Co-transfection of DNA and siRNA was performed using Lipofectamine 2000 and cells were harvested for analyses 60-72h post-transfection.

For total protein collection, cells were washed twice in cold PBS, scraped, and incubated in lysis buffer [50mM Tris-HCl (pH 7.4), 75mM NaCl, 0.5mM EDTA, 0.5% Triton-X, 1X protease inhibitor cocktail (Catalogue 4693159001, Sigma Aldrich), 1X phosphatase inhibitor cocktail (Catalogue 4906837001, Sigma Aldrich)] at 4°C for 20 min with moderate tumbling. Lysates were centrifuged for 5000 x g for 10 min and supernatants were stored at -20^ᵒ^C. For total RNA collection, 1mL TRIzol reagent (Catalogue 15596-018, Thermo Fisher Scientific) was added directly to cells following two PBS washes. For experiments involving LRRK2 kinase inhibition, cells were treated with 1 μM of DMSO-dissolved LRRK2 inhibitor GSK2578215A (Catalogue 4629, Tocris), or 2 μM LRRK2 inhibitor HG-10-102-01 (Catalogue 438195, EMD Millipore) for 3 hours immediately before cell harvest. Control cells were simultaneously treated with equivalent amounts of DMSO vehicle.

### Mice and tissue homogenization

C57BL/6 *HuD^+/-^* mice^48^ were previously generated by Hideyuki Okano (Keio University, Tokyo). Mice with the LRRK2 G2019S mutation knocked into the endogenous loci mice were a kind gift of Novartis^70^. Experimental mice were bred in two groups C57BL/6 *HuD^+/-^* x *LRRK2^WT/WT^* were crossed with C57BL/6 *HuD^+/-^* x *LRRK2^WT/WT^* to generate C57BL/6 *HuD^-/-^* x *LRRK2^WT/WT^* and C57BL/6 *HuD^+/+^* x *LRRK2^WT/WT^*. C57BL/6 *HuD^+/-^* x *LRRK2^G2019S/G2019S^* were crossed with C57BL/6 *HuD^+/-^* x *LrrkG^G2019S/G2109S^* to generate C57BL/6 *HuD^-/-^* x *LRRK2^G2019S/G2019S^* and C57BL/6 *HuD^+/+^* x *LRRK2^G2019S/G2019S^*. Mice were randomized into cages at weaning. At approximately 4 weeks of age, cortex, ventral midbrain, dorsal midbrain and striatum tissues were harvested. Tissues were flash frozen and stored at -80°C. For protein analysis, a fraction of each brain region was re- suspended in homogenization buffer [50mM Tris-HCl (pH 7.4), 140mM NaCl, 1% SDS, 1% Triton-X and 1X protease inhibitor cocktail] in 2.0mL tubes containing stainless steel beads. Tubes were placed in a MagNA lyser for two rounds of rapid oscillation at 7000rpm for 15 sec. Tissues were then incubated with gentle shaking at 4°C for 20 min and centrifuged twice at 10,000 x g for 10 min. Tissue lysates were stored at -80°C until further use. For total RNA analysis, fractions of brain tissue were resuspended in 1mL TRIzol reagent. For HuD-RNA immunoprecipitation assays, frozen midbrain tissues were crushed on dry ice using a steel Plattner’s Mortar and Pestle (Thomas Scientific) prior to incubation in IP lysis buffer [50mM Tris (pH 7.4), 150mM NaCl, 1% Nonidet P-40, and 1X protease inhibitor cocktail] with gentle tumbling for 30min at 4°C.

### Western blotting

Protein concentration was estimated using the Bradford Protein Assay (Catalogue 500-0122, Biorad). Lysates were diluted in Laemmli loading buffer [50mM Tris (pH 6.8), 2% SDS, 10% glycerol, 5% β-mercaptoethanol, 0.1M DTT] denatured at 99°C for 8-10 min and separated by 10% polyacrylamide gel electrophoresis (SDS-PAGE). Proteins were transferred to polyvinylidene fluoride (PVDF) membranes (Catalogue IPVH00010 , Fisher Scientific) and membranes were blocked in 5% milk for 1 hour at room temperature. Proteins were visualized using Luminata Crescendo Western HRP chemiluminescent substrate (Catalogue WBLUR0500, Fisher Scientific) to the membrane on an Image Quant LAS4010 Biomolecular Imager (GE Healthcare). Quantitative analyses on band intensity levels were performed using the Histogram function on Photoshop CS6 software with an area of fixed size. Mean band intensity was normalized to Tubulin or Actin levels.

### Enzyme-Linked Immunosorbent Assay (ELISA)

Seventy-two hours after transfection cell culture media was centrifuged at 1500 x g for 10 min and supernatants were immediately used in the Quantikine Human BDNF Immunoassay (Catalogue DBD00, R&D Systems). Values of a media blank were subtracted before calculating BDNF concentration based on a standard curve.

### Gel-shift Assay

10 pmol of RNA probes were 5’-labelled with 25 pmol of [γ-^32^P]ATP (3,000Ci/mmol, 10 mCi/ml, PerkinElmer) and T4 Polynucleotide Kinase (ThermoFisher) at 37°C. Probes were cleaned and unincorporated nucleotides removed through G25 Sephadex columns. Phosphorylation assays were performed in 25 μl as described above with 1.5 μg of GST-HuD or GST, 60 ng of wild-type, G2019S or D1994A LRRK2 recombinant proteins, 10 μM of ATP, in phosphorylation buffer, in absence or presence of kinase inhibitors (1 μM GSK2578215A or 2 μM HD). Binding assays were carried out at room temperature for 30 min with 2 μl of phosphorylation assays and 20,000 cpm of RNA probes in binding buffer (20 mM Tris pH 7.4, 1 mM EDTA, 100 mM KCl, 1 mM DTT, 5% Glycerol) in a total volume of 20 μl. At the end of the incubation, 2.5 μl of 10X loading buffer (100 mM Tris pH 7.4, 10 mM EDTA, 50% Glycerol, 0.1 % Xylene Cyanol, 0.1 % Bromophenol Blue) were added to the binding reaction and 10 μl were loaded and separated on a 4% acrylamide:bisacrylamide (29:1) gel. After electrophoresis, gels were dried on 3MM Whatman papers and autoradiographed using X-ray films.

### Immunoprecipitation

For RNA immunoprecipitation, IP lysis buffer also contained 40units/mL of RNaseOUT Recombinant Ribonuclease Inhibitor (Catalogue 10777019, Thermo Fisher Scientific). To pre-clear cell lysates, 10uL of washed Dynabeads® Protein-G magnetic beads (Catalogue 10004D, Thermo Fisher Scientific) were added to each lysate and discarded 20 min later. To immunoprecipitate the protein-complexes, 2 ug of antibody to the protein of interest or 2 ug of mouse IgG control antibody (eBioscience) was added to 800ug-900ug of protein lysate and incubated at 4°C for 2 hours. To capture the antibody-protein complexes, 20uL of magnetic beads were washed and then added to each IP. After 1 h samples were washed (5 min) with increasing NaCl concentration for each wash (150mM, 175mM and 200mM). For RNA-protein immunoprecipitations, 10% of the beads were resuspended in 20uL IP lysis buffer and Laemmli buffer for western blot, while remaining beads were resuspended in TRIzol reagent.

### Phosphorylation assay

GST-tagged mouse HuD or GST control proteins were a gift from Dr. Jocelyn Coté. Wild-type LRRK2, recombinant G2019S LRRK2 (aa. 970-2527) and recombinant D1994A LRRK2 (aa. 970-2527) human proteins were obtained from Thermo Fisher Scientific (A15197, PV4881 and PV6051). *In vitro* phosphorylation assays were carried out in a 25 uL volume by combining 1.5 ug of GST-HuD (or GST control) with 60 ng of wild-type, G2019S or D1994A LRRK2 in phosphorylation buffer (20mM HEPES (pH 7.5), 10mM MgCl2, 1mM EGTA, 2mM DTT, 1% DMSO and 1X PhosSTOP^TM^ phosphatase inhibitor cocktail). To initiate phosphorylation, 10μM ATP and 10μCi [γ-^32^P]ATP (Catalogue BLU002A500UC, Perkin Elmer) was added to each tube and reactions were incubated at 30°C for 1hr. Reactions were performed in parallel but without [γ-^32^P]ATP to confirm equal amounts of HuD and LRRK2 in each reaction by Western blot. Reactions were terminated by the addition of 1X Laemmli loading buffer and were subsequently boiled at 99°C for 5 min. Membranes were dried and exposed to autoradiography film for 24-72 hours before imaging.

### PhosTag

Phos-Tag SDS-PAGE gels consisted of a 12% acrylamide resolving gel containing 50 μM Phos-tag Acrylamide (Catalogue AAL-107S1, Wacko) and 100 μM ZnCl2. Samples were run and transferred using Western blot protocol above.

### Splicing RT-PCR

cDNA synthesis was performed using the iScript cDNA synthesis kit (Catalogue 170-8891, BioRad) and PCR using Taq DNA polymerase (Catalogue HTD0078, Biobasic). The cycling conditions were as follows: 95°C, 3min, [95°C for 30 sec, 55°C for 30 sec, 72°C for 30 sec] x 35 cycles, 72°C for 5 min. Samples were run on a 2% Agarose gel.

### Protein Identification by LC-MS/MS

Samples were prepared in a sterile work environment and separated on 4-20% Mini-PROTEAN TGX Pre-cast Protein Gels (Catalogue 165-3366, BioRad). Proteins were visualized using the PlusOne Silver Staining Kit (Catalogue, 17-1150-01, GE Healthcare Life Sciences), excised from gels and stored in 1% acetic acid. Proteins were digested in-gel using trypsin (Promega). Peptide extracts were concentrated by Vacufuge (Eppendorf). LC-MS/MS was performed using a Dionex Ultimate 3000 RLSC nano HPLC (Thermo Scientific) and Orbitrap Fusion Lumos mass spectrometer (Thermo Scientific) by the Ottawa Hospital Proteomics Core Facility. MASCOT software version 2.6 (Matrix Science, UK) was used to infer peptide and protein identities from the mass spectra. The observed spectra were matched against mouse sequences from SwissProt (version 2016-09) and also against an in-house database of common contaminants. The results were exported to Scaffold PTM Viewer (Proteome Software, USA) for analysis.

### RNA Extraction and RT-qPCR

Total RNA from cells and tissues was extracted with TRIzol reagent following the manufacturer’s instructions (Catalogue M0253L, Thermo Fisher Scientific). RNA was reverse transcribed using M-MuLV reverse transcriptase (New England Biolabs) supplemented with Oligo(dT)18 and dNTP mix (Catalogue R0192, Thermo Fisher Scientific) following the manufacturer’s protocol. qPCR reactions were carried out using GoTaq ® qPCR master mix (Catalogue A6002, Promega). Cycling conditions for all primer sets was as follows: 95°C for 2 min, (95°C for 15 sec, 60°C for 40 sec) x 40 cycles. Fold changes in mRNA levels were calculated using the ΔΔCt method. For SH-SY5Y experiments, mRNA levels were normalized to *GAPDH* and *TBP* mRNA, while in mice *Actin* was used for normalization.

### Immunoprecipitated RNA

cDNA was prepared from 7 uL of RNA using iScript Reverse Transcription Supermix (Catalogue 1708840, BioRad) according to manufacturer’s instructions. qPCR reactions were carried out using SsoAdvanced Universal SYBR Green Supermix (Catalogue 1725271, BioRad). Cycling conditions for all primer sets was as follows: 95°C for 3 min, (95°C for 10 sec, 60°C for 20 sec, 72°C for 20 sec) x 40 cycles. Technical replicates that produced undetectable Ct values or multiple/incorrect melt curve peaks were omitted from analysis. mRNA levels in HuD immunoprecipitations were normalized to levels in control IgG samples and expressed as fold change relative to the IgG. In experiments comparing mRNA enrichment in different conditions, mRNA quantities were normalize to the amount of HuD immunoprecipitated (using band intensity from western blots.)

### Microarray

Total RNA and RNA extracted from HuD immunoprecipitates from the midbrains of 4 week old mice was sent to the Centre for Applied Genomics at Sick Kids Hospital (Toronto, Canada) for microarray analysis. For immunoprecipitated RNA, equal volumes of RNA was reversed transcribed into cDNA. For total cellular RNA, 8000 pg RNA was reverse transcribed into cDNA. All DNA was prepared using the Affymetrix GeneChip™ Whole Transcriptome (WT) Pico Kit (Thermo Fisher Scientific) and labelled with biotin allonamide triphosphate. Labelled target DNA was hybridized for 16-18h at 45°C to Clariom™ D Mouse Pico GeneChip™ (Thermo Fisher Scientific). GeneChip assays were scanned using an Affymetrix GeneChip Scanner 3000.

### Bioinformatics analysis

Analysis of microarray data was performed using the Affymetrix Transcriptome Analysis suite v 4.0.1, and MTA 1.0 r3 annotations by Gareth Palidwor at the Ottawa Bioinformatics Core. Fold change analysis was performed using RMA-normalized (Robust Multi-array Average) conditions for both immunoprecipitated RNA and total RNA samples. The g:Profiler tool [https://biit.cs.ut.ee/gprofiler/] was used for gene ontology analysis^78^.

The density of neuronal ELAV binding motifs in 3’UTRs was quantified by extracting the 3’UTR from all known mRNAs detected in the dataset. Then, the number of neuronal ELAV binding motifs (UUU*UUU)^33^ was quantified per mRNA and normalized to the total 3’UTR length of the given mRNA.

### Site-directed mutagenesis

The human FLAG-HA-HuD plasmid was used to create mutations in the sites corresponding to mouse T149 (T144A), T174 (T169A) and both T149/T174 (T144A/T169A). Mutations were created with primers in the Reagents section below. Primers were designed to anneal ’back-to-back’ on opposing DNA strands in a non-overlap extension site-directed mutagenesis using Q5 High-fidelity DNA polymerase (Catalogue M0491L, New England Biolabs).

### Dual Luciferase Assays

A 216 nt fragment of the mouse BDNF 3’UTR including the HuD binding site (Allen et al. 2013), was cloned into the XbaI and NotI sites of the Renilla luciferase pRL-TK vector. pGL3.14 Firefly luciferase vector was co-transfected in U2OS cells to normalize for effects on transfection and gene expression. pFRT-DestFLAGHA- HuD was co-transfected where indicated. Co-transfection of DNA and siRNA was performed with Lipofectamine 2000. Cells were harvested 48 hours post-transfection and lysed in passive lysis buffer and analyzed using the Promega Dual Luciferase Stop and Glo Kit. Data were normalized to cells expressing empty pRL-TK vector transfected with control siRNA.

### Gateway cloning/Protein purification

Wild-type and mutant pFRT-DestFLAGHA-HuD plasmids were used to produce GST-HuD and GST-HuD (T149A, T174A) bacterial expression plasmids using Gateway cloning. pENTR4-ELAVL2 (Addgene #65752) and pENTR4-ELAVL3 (Addgene #65752) were used as Entry plasmids for Gateway cloning to produce GST- HuB and GST-HuC, respectively. Plasmids were transformed in BL21 (DE3) competent *E.coli*. An overnight culture was diluted 1 in 100 in LB broth including ampicillin and protein expression was induced at OD600 0.4 with 0.2 mM Isopropyl β-D-1-thiogalactopyranoside for four hours at 30 degrees Celsius. Bacterial culture was centrifuged at 10,000 x 30 min (4°C) and lysed with 1 mg/mL Lysozyme in resuspension buffer (50mM Tris, pH 8, 200 mM NaCl, 1mM EDTA, 1mM DTT, 1% Triton, 5% glycerol and protease inhibitor tablet). Lysate was sonicated and centrifuged at 12,000xg for 30 min. Supernatant was loaded on a Glutathione-agarose column and eluted with 10 mM reduced glutathione in resuspension buffer.

### Behaviour and Motor Co-ordination Analysis

Behaviour analysis was completed in a dark light cycle to which mice were acclimated for 2 weeks prior to testing. Mice were acclimated to the test room for 30-60 min prior to testing. Mice were tested at 10 weeks of age. Mice were randomized into group cages at weaning based on randomized numbered ear tags by independent personnel (MTT, AS) to enable blinded testing by ON and personnel of the University of Ottawa Behaviour Core Facility.

#### Beam Break

General locomotor activity was assessed using Beam Breaks in a novel cage during the dark cycle. Mice were individually housed and tested for a four-hour period. Ambulatory activity was included in analysis using Fusion software (Omnitech Electronics).

#### DigiGait

Gait analysis was tested using DigiGait. Mice were individually placed on the belt with the speed of 18cm/sec at an 8 degree incline. Video data lasting at least three seconds was used in the analysis, pooling data for right and left limbs for each mouse. Analysis showed no significant difference between left and right limbs in any group.

## Statistical Analysis

Statistical analysis was performed with GraphPad Prism. Two-tailed student’s t-tests were performed in cases comparing two independent groups of data. For data with more than two independent groups, a one-way ANOVA (analysis of variance) test was performed with a Tukey post-hoc test. Hypergeometric tests were performed to identify statistical significance in microarray experiments. Statistical significance was represented with the following notation: *p<0.05; **p<0.01, ***p<0.001, etc.

### Reagents

#### siRNAs

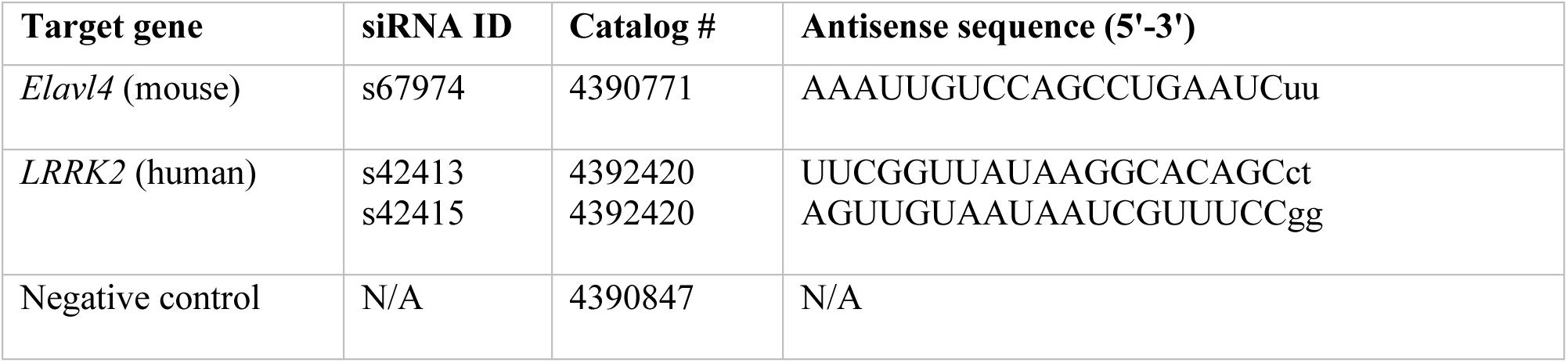

#### Antibodies

| Antibody | Dilution | Supplier | Catalogue # |
| --- | --- | --- | --- |
| **Primary** |  |  |  |
| α-synuclein | 1:5000 | Cell Signalling Technology | 2628 |
| α-Tubulin | 1:20,000 | Thermo Fisher Scientific | PA5-22060 |
| β-Actin | 1:10,000 | Sigma | A5441-.2ML |
| BDNF | 1:3000 | Abcam | Ab205067 |
| eIF4A (C32B4) | 1:10,000 | Cell Signalling Technology | 2013S |
| FLAG (M2) | 1:5000 | Sigma | F1804-IMG |
| HuD (H-9) | 1:5000 | Santa Cruz | sc48421 |
| HuD | 1:3000 | Abcam | Ab96474 |
| LRRK2 (MJFF2[c41-2]) | 1:1000 | Abcam | Ab133474 |
| LRRK2 P<sup>Ser935</sup> (UDD2-10(12)) | 1:1000 | Abcam | Ab13340 |
| P-Thr (H2) | 1:1000 | Santa Cruz | sc5267 |
| p21 (EPR362) | 1:3000 | Abcam | Ab109520 |
| **HRP-secondary** |  |  |  |
| Goat anti-mouse | 1:5000 | Jackson ImmunoResearch | 115-035-174 |
| Goat anti-rabbit | 1:5000 | Jackson ImmunoResearch | 111-035-144 |
| **Isotype control (for immunoprecipitation)** |  |  |  |
| Mouse IgG1 K immunoglobulin | N/A | eBioscience | 14-4714-82 |
| Goat anti-rabbit | N/A | Jackson ImmunoResearch | 111-035-144 |

#### Primers

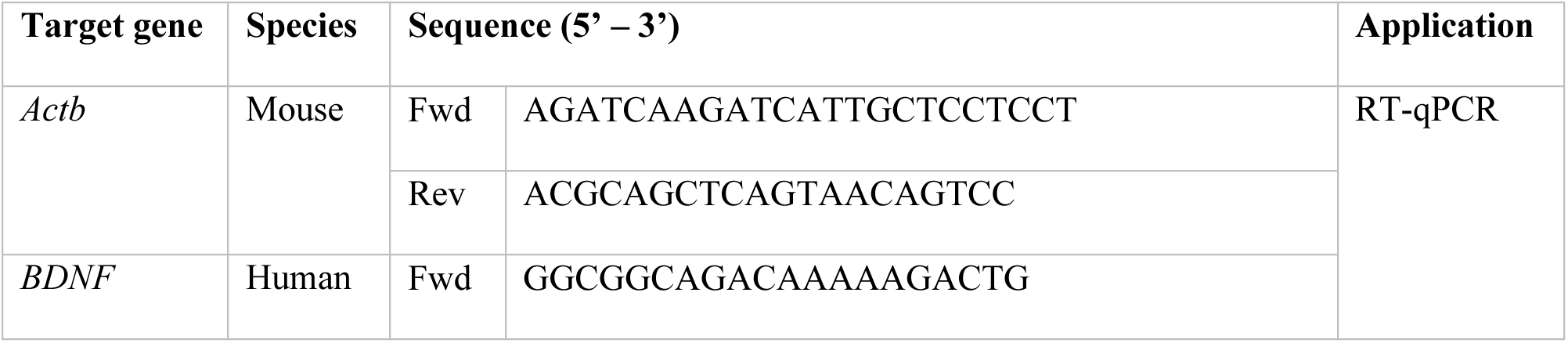
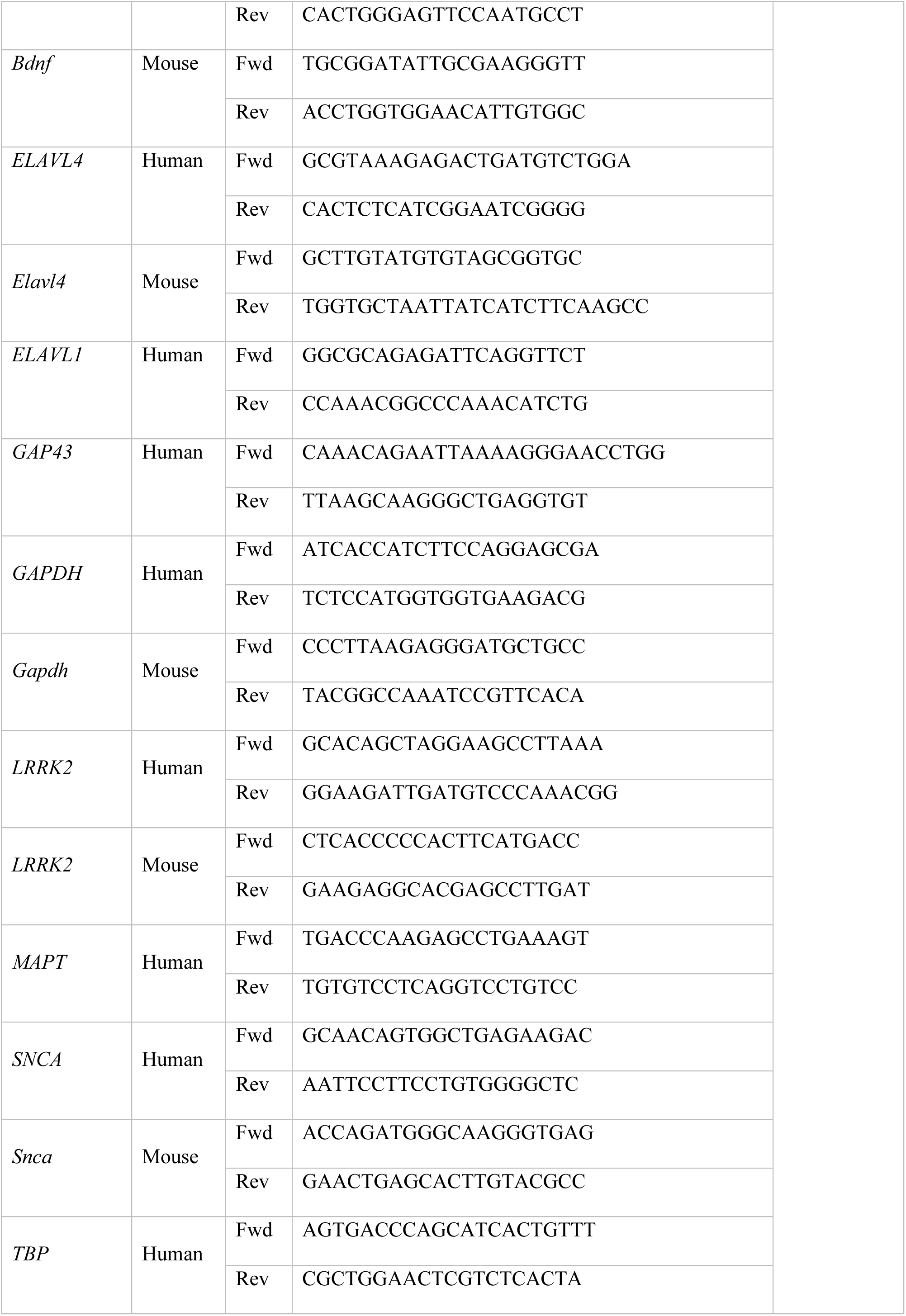
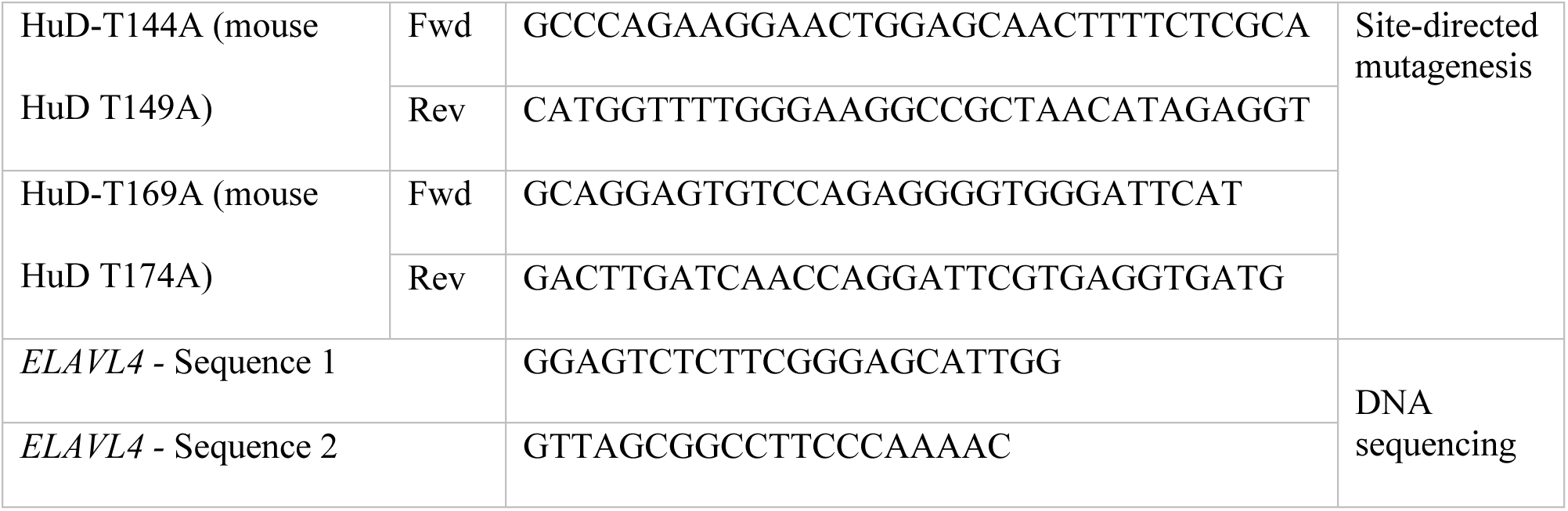

